# The nucleolus is a mechanosensitive condensate that adapts ribosome biogenesis to mechanical forces

**DOI:** 10.64898/2026.07.02.731374

**Authors:** Yuthika Shetty, Kenny O. Elias, Sara Badawi, Lydia Pernet, Anne-Sophie Ribba, Christiane Oddou, Ludivine Wacheul, Lucid Belmudes, Eve Moutaux, Christiane Zorbas, Sandrine Fraboulet, Yohann Coute, Fabian Erdel, Denis L.J. Lafontaine, Monika E. Dolega

## Abstract

Intracellular compartmentalization is fundamental to cellular organization, yet mechanobiology has been largely understood through membrane-delimited structures and associated signaling pathways. Whether mechanical forces directly regulate biomolecular condensates, which organize many core cellular functions, remains largely unknown. This question is particularly relevant for the nucleolus, a prominent nuclear condensate that coordinates ribosome biogenesis and is known to remodel in response to diverse biochemical perturbations, placing it at the interface between cellular state and biosynthetic control. Here, we show that mechanical compression remodels nucleolar organization and reduces (ribosomal DNA) rDNA transcription, and identify nucleolin as a key mediator of this adaptive response. Compression induces rapid and reversible redistribution of nucleolin from the nucleolus to the nucleoplasm, accompanied by reduced occupancy at rDNA promoter regions and changes in rDNA transcription and precursor rRNA processing. The nucleolar response occurs independently of classical post-translational regulation of nucleolin and instead depends on the rate of nuclear deformation, with nucleolar organization and function scaling with nuclear volume loss, supporting a mechanism of biophysical regulation. Together, our findings establish the nucleolus as a mechanosensitive condensate and reveal dual regulation of ribosome biogenesis by mechanical compression, through rapid nucleolin-based biophysical adaptation followed by slower epigenetic remodeling.

## INTRODUCTION

Epithelial tissues constantly experience mechanical forces arising from growth, crowding, stretch, and confinement. These forces are increasingly recognized as key regulators of tissue homeostasis, shaping proliferation, differentiation, and survival decisions that maintain tissue architecture [1,2]. Mechanical inputs are transmitted through the cytoskeleton and the LINC complex to the nucleus, where they influence nuclear architecture, chromatin organization, and gene expression programs [3,4]. Yet, despite growing appreciation that many nuclear functions are spatially organized within biomolecular condensates, whether mechanical cues directly regulate condensate organization and activity remains largely unexplored. This question is particularly relevant for the nucleolus, a prominent nuclear condensate that governs ribosome biogenesis, a major biosynthetic process that must be dynamically tuned to cellular and tissue demands. Recent studies suggest that rDNA transcription can be influenced by the cellular microenvironment [5–7], but the mechanisms linking mechanical signals to nucleolar function remain unknown.

The nucleolus is the largest membraneless organelle of the nucleus and the site where rDNA transcription, ribosomal RNA (rRNA) processing, and early ribosomal subunit assembly are coordinated. These processes occur within a spatially organized multilayered architecture composed of distinct functional subcompartments - the fibrillar center (FC), dense fibrillar component (DFC), and granular component (GC) - that correspond to successive steps of ribosome production [8–11]. High-end fluorescence microscopy has revealed additional nucleolar layers, including the periphery of DFC (pDFC) and nucleolar rim [9,12]; additional subphases are likely to be discovered [13]. Beyond its role in ribosome assembly, the nucleolus also functions as a central hub for cellular stress responses, adjusting ribosome biogenesis in response to diverse perturbations [14,15]. Under stress such as genotoxic, heat shock, oxidative and osmotic stresses, the nucleolus can undergo a range of structural and compositional changes, including disruption of its ultrastructure [14], transcriptional repression of rDNA [16], sequestration of misfolded proteins [17,18], and/or redistribution of core nucleolar factors. Proteins such as fibrillarin [19], nucleophosmin [20], UBF [21], and nucleolin [19,22] exhibit stress-dependent changes in localization and interactions that accompany the reorganization of nucleolar architecture and regulation of ribosome biogenesis. Despite extensive characterization of nucleolar stress responses, how mechanical cues might engage nucleolar regulatory pathways remains unknown. This gap is conceptually intriguing because the nucleolus lacks a surrounding membrane and is thought to assemble through liquid–liquid phase separation [23], making its responsiveness to mechanical signals non-trivial from the perspective of classical membrane-based mechanobiology.

The formation and stability of the nucleolus are governed by multivalent interactions between nucleolar proteins and rRNA that drive liquid–liquid phase separation [23]. The material properties and organization of such condensates are highly sensitive to their physicochemical environment, including macromolecular concentration, ionic strength, and the degree of macromolecular crowding [24]. Mechanical perturbations can profoundly alter these parameters by inducing changes in cell and nuclear volume, intracellular ion balance, and the volume fraction occupied by macromolecules. Such shifts are therefore expected to directly influence the phase behavior and internal organization of nucleolar condensates [25]. However, whether mechanical cues regulate nucleolar organization and ribosome biogenesis through these physicochemical mechanisms remains largely unexplored.

Here we investigate the mechanisms underlying nucleolar adaptation to mechanical stress. Using multiple cellular compression modalities to probe nucleolar responses to mechanical load, we show that mechanical compression induces rapid remodeling of nucleolar organization accompanied by reduced rDNA transcription. We identify nucleolin as a key mediator of this adaptive response, undergoing rapid and reversible translocation from the nucleolus to the nucleoplasm under compression. Nucleolin redistribution is associated with its depletion from rDNA promoter regions and with reduced rDNA transcription and rRNA processing. Notably, nucleolin translocation occurs without detectable post-translational modifications and correlates with nuclear deformation and volume loss under compression, supporting a mechanism driven by changes in the physicochemical environment of the nucleus. Together, these findings establish the nucleolus as a mechanosensitive biomolecular condensate and identify a mechanism by which mechanical compression directly modulates ribosome biogenesis.

## RESULTS

### Mechanical compression remodels nucleolar ultrastructure and reduces rDNA transcription

In our previous work, we showed by RNA-seq that 4hours of extrinsic mechanical compression induces differential expression of ribosomal proteins and snoRNA transcripts in an MDCK epithelial model [26], suggesting potential regulatory role of mechanical stress in ribosome biogenesis. Building on this, we investigated how mechanical forces reorganize nucleolar architecture in MCF10A cells, a genetically well-characterized, non-transformed, human epithelial model [27]. Mechanical compression was achieved through the cultivation of cells for 48 hours on pre-stretched PDMS membranes followed by instantaneous release of stretch, imposing 30% biaxial strain on the MCF10A monolayers as described previously in the literature [28] (Figure. 1A). This release generates in-plane compression of the epithelial sheet, increasing monolayer density and recapitulating mechanical constraints associated with epithelial crowding during tissue growth or confinement. We observed that this in-plane compression triggered nucleolar reorganization. Mean nucleolar area increased significantly (Figure 1B) and was accompanied by a higher proportion of nuclei containing a single large nucleolus suggesting increased frequency of fusion of individual nucleoli (Figure 1C). These changes were evident 4 hours after compression, coinciding with previously reported transcriptional changes in genes associated with ribosome biogenesis[26].

**Figure 1.**
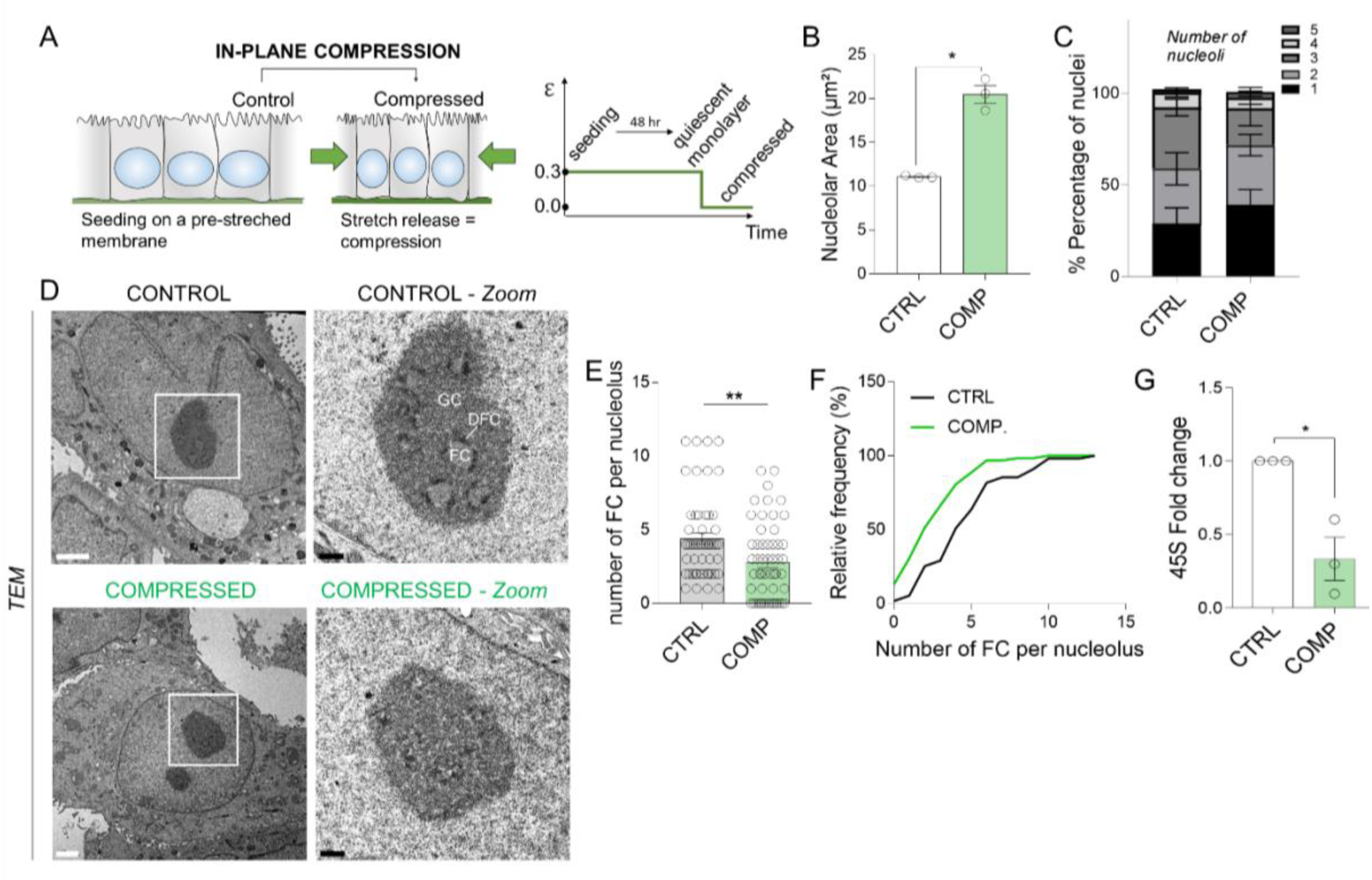
Mechanical compression remodels nucleolar ultrastructure and reduces rDNA transcription. A) Schematic representation of the in plane mechanical compression and its dynamics by a strain profile over time. B) Nucleolar area increases in cells exposed to in-plane compression for 4 hours (N=3 independent experiments with n>50 per condition/experiment; *p= 0.0115 in unpaired t-test with Welch’s corr.). C) Quantification of nucleolar number per nucleus in control and compressed monolayers at 4 hours (N=5 independent experiments with n>50 per experiment per condition). D) Representative TEM images of nucleoli in control and compressed cells. Zoomed images highlight nucleolar substructures and loss of fibrillar centers in compressed cells. Fibrillar centers (FC), dense fibrillar component (DFC), and granular component (GC) are indicated in the control image. (Scale bar is 2µm in control, compressed and 0.5 µm for the zoomed images). E) Number of FCs per nucleolus quantified in control versus mechanically compressed cells (N=3 independent experiments with n>15 per condition/experiment; p=0.0014 in Mann Whitney test). F) Relative frequency distribution of FC counts per nucleolus in all experiments shows a marked decrease in number of FCs in compressed cells compared to control. G) RT-qPCR analysis of 45S rRNA levels showing reduced relative abundance in compressed cells (N=3 independent experiments; p=0.0105 in the unpaired t-test).

To further examine whether compression-induced changes in nucleolar morphology were reflected at the ultrastructural level, we performed transmission electron microscopy (TEM). Ultrastructural analysis revealed a clear reorganization of nucleolar architecture under compression (Figure 1D). Compressed cells displayed significantly fewer fibrillar centers (FCs) per nucleolus, which were more difficult to distinguish as compared to uncompressed controls (Figure 1E, 1F). Because FCs are closely associated with active rDNA repeats and organize Pol I transcription complexes at the FC–DFC interface [10], their reduction suggested mechanical forces actively remodeled the transcriptional output within the nucleoli. RT-qPCR experiments on compressed cells further confirmed this hypothesis, showing a significant reduction in nascent 45S pre-rRNA, measured using primers targeting the 5′ external transcribed spacer (5′ETS) (Figure 1G). Together, these data indicate that mechanically driven nucleolar reorganization represents an adaptive response to mechanical stress rather than a complete transcriptional shutdown, in contrast to the known effect of RNA polymerase I inhibition (e.g. Actinomycin D treatment), which abolishes rDNA transcription and leads to nucleolar segregation and disassembly [21].

Because nucleolar function is spatially organized across distinct internal compartments and redistribution of nucleolar factors is a hallmark of canonical nucleolar stress responses, we next examined whether mechanical compression alters the distribution of nucleolar proteins. To this end, we selected representative nucleolar markers spanning the major nucleolar subcompartments (schematically illustrated in Figure 2A) - upstream binding factor (UBF; FC), fibrillarin (DFC), nucleolin (C23/NCL; GC, pDFC, DFC), and nucleophosmin (B23/NPM; GC) - which have been previously described to relocalize under nucleolar stress conditions. Immunofluorescence analysis revealed that, among the nucleolar markers examined, only nucleolin redistributed from the nucleolus to the nucleoplasm upon compression (Figure 2B). Nucleolin normally spans multiple nucleolar regions, including the DFC, its peripheral region (pDFC), and the GC. Given its presence across these compartments, nucleolin displacement could potentially influence ribosome biogenesis at several stages of the process. Interestingly, western blot analysis showed that total cellular levels of fibrillarin, UBF, NPM, and NCL remained largely unchanged under compression (Figure 2C), emphasizing that the response primarily reflects protein relocalization rather than strong changes in overall abundance.

**Figure 2.**
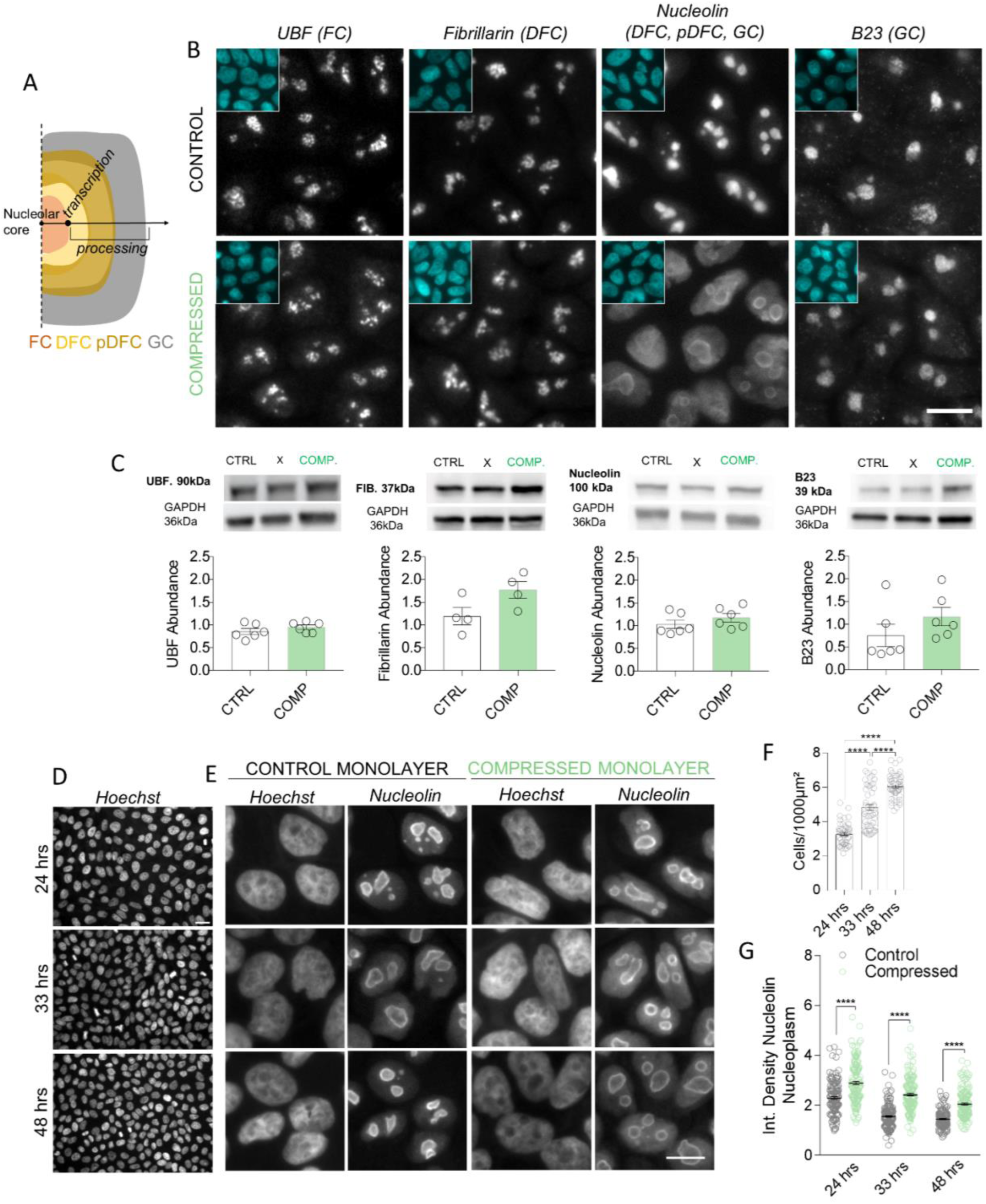
Nucleolar remodeling under mechanical compression is accompanied by nucleolin redistribution to the nucleoplasm. A) Schematic representation of nucleolar subcompartments, including the fibrillar center (FC), dense fibrillar component (DFC) with its peripheral region (pDFC), and the granular component (GC). Major nucleolar activities, including rDNA transcription and rRNA processing, are indicated to illustrate their spatial organization relative to the markers shown in panel B. B) Representative images of control and compressed cells stained for nucleolar proteins shows a strong change in the localization of nucleolin under compression. Each panel includes a cyan inset showing Hoechst-stained nuclei (N>4 of independent experiments). C) Total abundance of nucleolar proteins fibrillarin, UBF, B23 and Nucleolin does not change under compression as shown by WB quantification. “X” indicates conditions not presented in this study. (N>4 of independent experiments for each condition; p=n.s. in Mann-Whitney test). D) Representative images of different monolayer densities used in the experiments. E) Representative images of Hoechst and nucleolin for control and compressed monolayers at different density time points. F) Quantification of cells/1000 µm2 of different monolayer densities used in the experiments. (N=3 independent experiments; data was pooled together; **** p<0.00001 in Mann Whitney test). G) Quantification of nucleolin levels in control and compressed conditions in monolayers of different densities shows increased accumulation of nucleolin in nucleoplasm under compression. (N=3 independent experiments, with total 200 cells analyzed across experiments; data was pooled together; **** p<0.00001 Mann Whitney test).

To determine whether this response is consistent across varying physiological mechanical states induced by different cell densities, we performed immunofluorescence staining for nucleolin on progressively denser monolayers of cells cultured for 24, 33, and 48 hours (Figure 2D, 2F). Across all density timepoints, mechanical compression induced significant nucleolin accumulation in the nucleoplasm (Figure 2E, 2G), suggesting a conserved, stress-induced response mechanism. These findings therefore establish nucleolin delocalization as a core component of the cellular response to externally applied mechanical stress.

### Mechanical compression increases nucleolin mobility and drives its rapid redistribution from the nucleolus

To visualize nucleolin translocation in real time, we performed live-cell microscopy using MCF10A cells stably expressing an N-terminally RFP-tagged nucleolin construct (*see Materials and methods*). Because the PDMS-based Flex-plate biaxial compression setup is not compatible with live imaging, we used a microscopy-compatible, pressure-controlled dynamic cell confiner, which imposes mechanical compression through controlled axial confinement of epithelial monolayers to a defined height [29] (Figure 3A). To confirm that axial confinement produces the same phenotype as in-plane compression, we performed antibody-based immunofluorescence staining in MCF10A wild type cells, which similarly revealed nucleolin accumulation in the nucleoplasm (Figure 3B). Consistently, western blot analysis showed that total nucleolin levels remain unchanged under compression (Figure S1A), indicating that the observed phenotype reflects protein relocalization rather than chandegradation.

**Figure 3.**
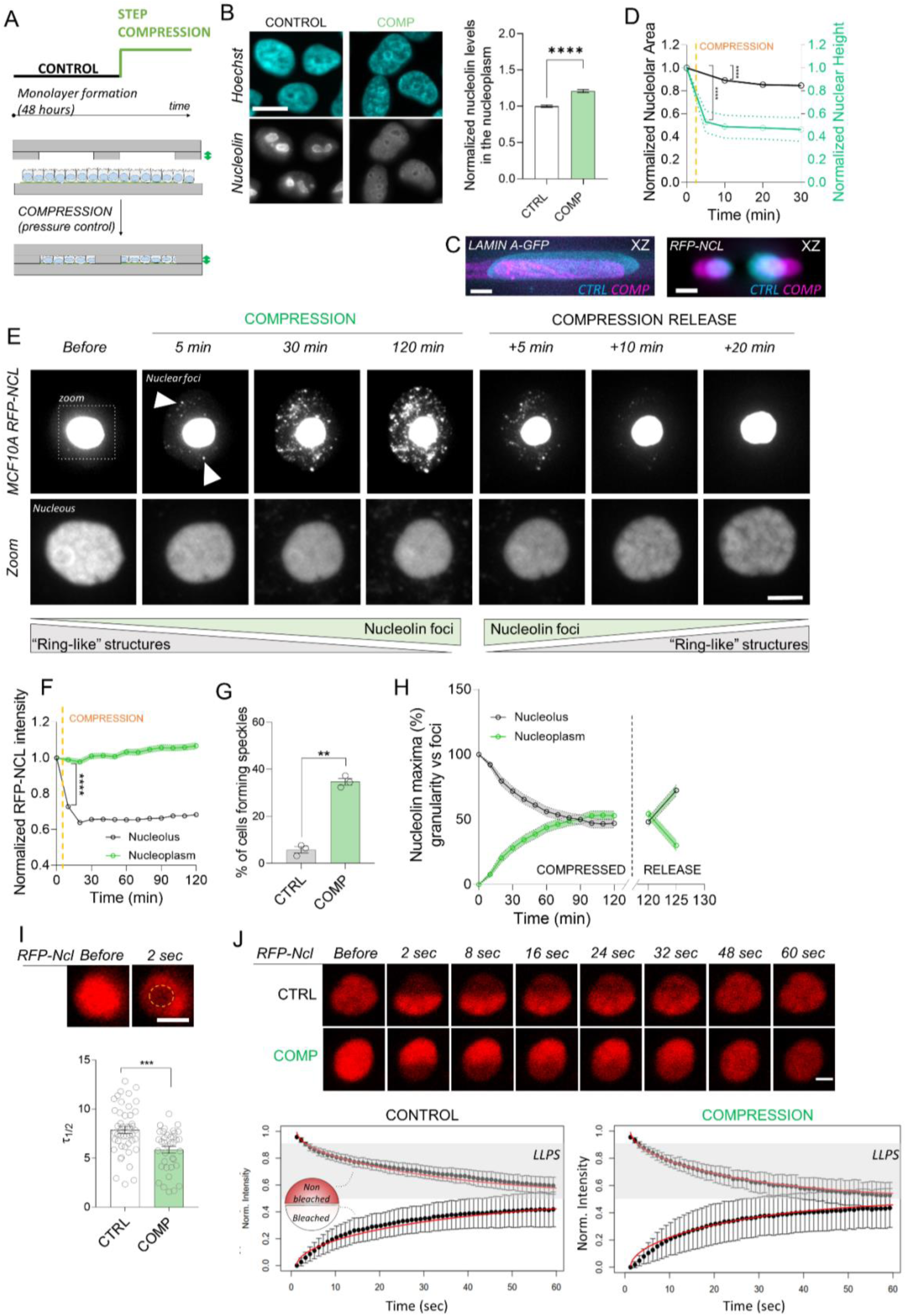
Mechanical compression increases nucleolin mobility and drives its rapid redistribution from the nucleolus. A) Schematic representation of axial mechanical compression (confinement) system and its dynamics represented by a strain profile over time. Deformation is regulated by the height of the PDMS pillars (green arrows). B) Representative images of MCF10A cells stained for nucleolin depicting nucleolin translocation to nucleoplasm under axial compression and nucleolin quantification of intensity in nucleoplasm (Scale bar=10µm, N=3 independent experiments, n=25 of cells per condition per experiment; **** p<0.0001 in Mann Whitney test). C) Representative images of Lamin chromobody-GFP and NCL-RFP showing nuclear and nucleolar deformation respectively before (cyan) and within 5 minutes after compression (Magenta) (Scale bar=2 µm). D) Normalized changes in nuclear height (green) and nucleolar area (black) upon compression show a 50% heigh and 10% deformation within 10 minutes of compression (pillar height= 4µm). (for nuclear heigh: N=3 independent experiments with n=5 nuclei analyzed per condition per experiment. For nucleolar area: N=3 independent experiments, with n=40 nucleoli analyzed per condition per experiment; **** p<0.0001 in Mann Whitney test. Each time point is represented by mean+SEM.) E) Representative time-lapse images of NCL-RFP expressing cells captured before and after axial compression and following compression release. Two sets of contrast-adjusted images of the same cell highlight nucleolar “granularity” changes and the emergence of nucleolin foci in nucleoplasm (Scale bar=2 µm). F) Quantified change in nucleolin (RFP-NCL) intensity within nucleolus and nucleoplasm under compression showed significant decrease of its intensity within nucleolus and a progressive increase in nucleoplasm. (N=3 independent experiments, with n=25 nuclei analyzed per condition per experiment; **** p<0.0001 in Mann Whitney test. Each time point is represented by mean+SEM.) G) Quantification of fluorescent nucleolin foci appearance in nucleoplasm revealed 37% of the subpopulation of cells with this phenotype. (N=3 independent experiments with n>87 cells per condition per experiment; **p<0.01 in Unpaired t test with Welch’s correction) H) Distribution of fluorescence maxima under compression within individual cells was analyzed in time. Increasing nucleolin foci accumulation in the nucleoplasm (green) with decreasing abundance within the nucleolar ultrastructure (gray) was observed under compression. Upon release partial recovery is observed within first 5 minutes (N=3 independent experiments with n=15 cells per condition per experiment. Each time point is represented by mean+SEM) I) Representative NCL-RFP images acquired immediately before FRAP and 2 s after bleaching within the nucleolus, with the bleached area marked in yellow. Quantification of half-time of recovery (τ₁/₂) revealed enhanced nucleolin mobility within the nucleolus under axial compression. (Scale bar=2 µm, N=3 independent experiments with n=15 nucleoli analyzed per condition per experiment; *** p<0.001 in paired t-test). J) Time-series images of NCL-RFP labeled nucleoli bleached in one half of their volume. Plots of intensity changes show mean normalized intensity in the bleached and non-bleached halves (illustrated in the schematic within the graph) revealed faster recovery kinetics within compressed cells (τ = 23 s) compared to controls (τ = 31 s). Red curves represent the three-parameter fit model indicated in the results section. (Scale bar=2 µm, N = 2 independent experiments with n = 5 nucleoli per condition per experiment).

Axial confinement also enabled real-time analysis of cellular deformation and nucleolin dynamics during compression. Upon confinement, both the nucleus and nucleolus deformed instantaneously, as visualized using a lamin chromobody and RFP-nucleolin (Figure 3C). Nuclear height decreased by approximately 50% immediately after confinement (Figure 3C,3D). The nucleolus also underwent vertical compression, accompanied by a reduction in projected area of ∼10%, reaching a maximum decrease of ∼15% after 30 minutes of compression (Figure 3D, Figure S1B). Together, these observations reveal a dynamic morphological adaptation of nucleoli to mechanical compression.

To characterize how nucleolin redistributes during mechanical compression, we monitored its nucleolar localization over time by live-cell imaging (Figure 3E). Nucleolin translocation was detectable as early as 5–10 minutes after compression; the earliest time interval accessible following microscope readjustment after initiating compression. Nucleolin exhibited a maximal nucleolar depletion of ∼38% at 30 minutes of compression (Figure 3F). Notably, time-lapse imaging showed that nucleolin is initially enriched in visually distinct circular regions (“ring-like” structures; *see Zoom in insets*), within the nucleolus as recently described by super resolution microscopy [9], giving the organelle a characteristic textured appearance. From these nucleolar subdomains, nucleolin gradually redistributes to the nucleoplasm, and in roughly 40% of cells it forms discrete fluorescent foci dispersed throughout the nucleoplasm over 2 hours of compression (Figure 3G). To characterize the dynamic displacement of nucleolin from nucleolar core to nucleoplasmic foci under compression, we quantified the spatial distribution of nucleolin fluorescence maxima within individual cells. The maxima of nucleolin fluorescence intensity progressively shifted from the nucleolus to the nucleoplasm, highlighting progressive re-localization of nucleolin to the nucleoplasm under mechanical compression (Figure 3H). Strikingly, this translocation was reversible, with nucleolin relocalizing to the nucleolus within ∼5 minutes after release of compression (Figure 3H, Figure S1C). This recovery was accompanied by the disappearance of nucleoplasmic nucleolin foci, the restoration of nucleolin fluorescence maxima within the nucleolus, and the reappearance of nucleolin ring-like texture in nucleolar regions. Together, these observations reveal a rapid and reversible bidirectional trafficking of nucleolin that is tightly regulated by mechanical compression.

Given the rapid redistribution of nucleolin under mechanical compression, we next examined its mobility within the nucleolus using fluorescence recovery after photobleaching (FRAP). The half-time of recovery averaged ∼7.9 s in uncompressed MCF10A cells, consistent with previously reported values [30]. Under compression, we observed a decrease to ∼5.5 s (Figure 3I), indicating increased nucleolin mobility within the nucleolus during mechanical compression. In contrast, nucleophosmin-GFP exhibited no significant changes in its recovery kinetics under confinement (Figure S1D), highlighting the specific impact of compressive force on nucleolin dynamics. Additionally, half-FRAP experiments were performed on RFP-nucleolin and nucleophosmin-GFP to probe the internal mixing dynamics and interfacial barrier strength of the nucleolus under mechanical confinement (Figure 3J, Figure S1E). The recovery curves were adequately captured by a three-parameter model) [31], with deviations falling within the experimental error. Consistent with the spot-FRAP measurements, half-FRAP analysis for RFP-NCL confirmed that the characteristic recovery time τ of nucleolin decreased under compression (τ = 23 s versus 31 s in controls), indicating increased intra-nucleolar mobility and mixing dynamics. The recovery times in half-FRAP are larger than those of spot-FRAP because the bleach regions are larger. The fit yielded low values of the permeability parameter h in both control and compressed cells (h = 7 × 10⁻⁴ and h = 1 × 10⁻⁴, respectively), indicating that nucleoli exhibit an interfacial barrier that attenuates fast diffusive exchange of nucleolin both under normal conditions and under compression. In contrast, half-FRAP analysis of nucleophosmin-GFP revealed similar characteristic recovery times in control and compressed cells (τ ≈ 9 s in both conditions), despite a modest increase in the immobile fraction under compression, while the interfacial barrier strength remained comparable between conditions (Figure S1E). Together, these findings indicate that mechanical compression enhances nucleolin mobility and exchange within the nucleolus, thereby promoting dynamic trafficking and mechanosensitive control of its subcellular localization.

### Nucleolin redistribution under mechanical compression is associated with altered ribosome biogenesis

To investigate the functional consequences of nucleolin translocation under axial confinement, we measured rDNA transcription using 5-EU Click-iT RNA imaging and RT–qPCR of 45S pre-rRNA in control and compressed cells. By quantifying the intensity of Click-iT probe within the nucleoli, we measured a significant reduction in nascent rRNA transcript levels (∼40% decrease) as early as 15 minutes post mechanical compression (Figure 4A), indicating acute downregulation of rDNA transcription. Upon release from confinement, RNA transcript levels within the nucleolus did not recover 15 minutes post release of compression, the timescale of nucleolin translocation back to the nucleolus (Figure 4A). However, discrete fluorescent foci indicative of transcriptional re-initiation were observed in a subpopulation of cells, consistent with the expected delay between Pol I reactivation and the accumulation of a detectable nucleolar EU signal [32]. RT-qPCR analysis of 45S pre-rRNA levels corroborated these findings, revealing significant reduction in 45S rRNA transcript abundance at 2 hours of sustained mechanical compression, followed by a mild increase 1hour post release of compression suggesting that the complete rate of recovery extends beyond this time point (Figure 4B). Notably, rDNA transcription levels showed greater variability during the recovery phase. These findings link compression-induced changes in nucleolin mobility, to the corresponding suppression and reactivation of rDNA transcription.

**Figure 4.**
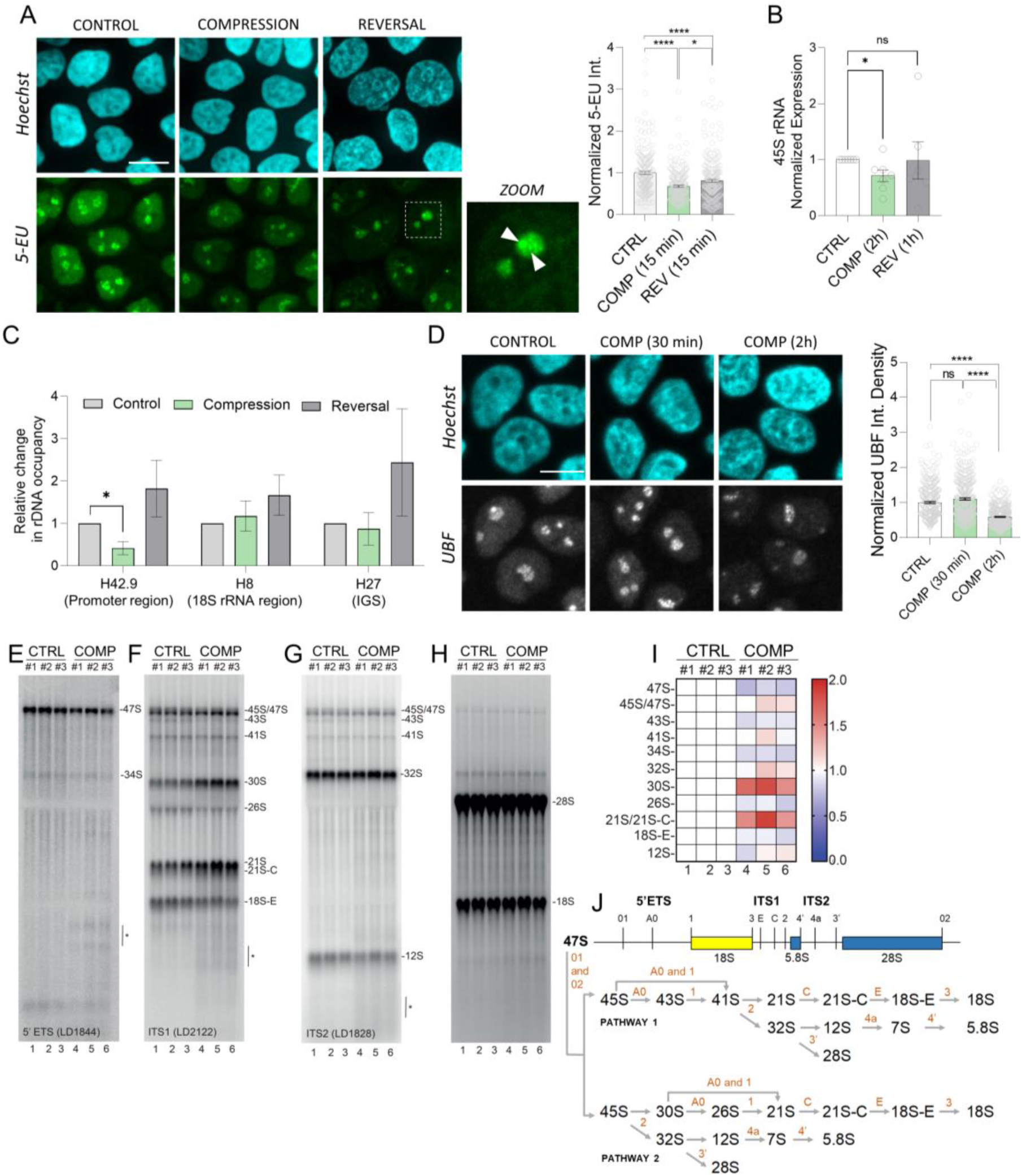
Nucleolin redistribution under mechanical compression is associated with altered ribosome biogenesis. A) Representative images of 5-EU Click IT assay and corresponding Hoechst images. Quantification showed marked reduction in nascent rRNA levels within the nucleolus 15 minutes post compression and 15 minutes post release. Zoom image shows bright fluorescent spots indicating rDNA reinitiation. (Scale bar=10 µm, N=3 independent experiments with at least 210 nucleoli per condition analyzed; Data are pooled together and presented as mean +/- SEM; *= 0.0118 and **** p<0.0001 Mann-Whitney test) B) RT-qPCR analysis of 45S rRNA levels showing reduced relative abundance in compressed cells and a variable trend of recovery upon reversal (N>6 independent experiments, * p=0.0105 in the unpaired t-test). C) Nucleolin ChIP shows significant decrease of nucleolin at the promoter region of rDNA (H42.9) at 2h of compression and no difference at the 28s rRNA region (H8) and intergenic sequence (H27). Nucleolin enrichment at rDNA promoter is restored 1h post release of compression. regions (N=3 independent experiments, * p=0.05 in the Mann Whitney test) D) Representative images of UBF (RNA pol I transcription factor) staining and Hoechst at 30 min and 2 h post-compression. Quantification revealed no change at 30 min but significant decrease of UBF within the nucleolus at 2 h. (Scale bar=10 µm, N=3 independent experiments with at least 338 nucleoli per condition; Data are pooled together and presented as mean +/- SEM; **** p<0.0001 in Mann-Whitney test) E-G) Northern blot probing with radioactively labelled oligonucleotides (Table S1). Membranes were hybridized with probes targeting the 5’ ETS (panel E), ITS1 (F), and ITS2 (G). (N=4 independent experiments, * indicates degraded fragments) H) Ethidium bromide staining to visualize large RNAs (18S, 28S). I) Heatmap representation of PhosphorImager quantification, showing reproducible changes across biological replicates showing rRNA-processing defects across biological replicates. J) Simplified schematics of the human pre-rRNA processing pathway, with cleavage sites and probe binding positions indicated.

To determine whether the compression-induced changes in rDNA transcription are associated with altered nucleolin engagement at rDNA loci, we next examined nucleolin occupancy across the rDNA repeat. Nucleolin is essential for rDNA transcription, where it promotes chromatin accessibility [33] and associates with the transcriptional machinery at rDNA promoter regions [34]. We therefore, performed nucleolin ChIP to assess how its occupancy is altered by compression across rDNA regions identified in previously published nucleolin ChIP-seq datasets [34]. Interestingly, ChIP analysis of compressed cells revealed a pronounced depletion of nucleolin at the promoter region (H42.9), while its occupancy at the 28S rDNA (H8) and intergenic spacer (H27) regions remained largely unchanged as compared to control (Figure 4C). rDNA promoter-associated nucleolin levels show a trend of recovery at 1 hour post release of compression, coinciding with nucleolin re-localization to the nucleolus and resumption of rDNA transcription. Moreover, immunofluorescence for the RNA Pol I transcription factor UBF shows its steady nucleolar enrichment at 30 minutes of compression, followed by a marked decrease at 2 hours (Figure 4D). The distinct temporal dynamics of rapid nucleolin displacement within 5 minutes and sustained UBF enrichment at 30 minutes suggesting continued RNA Pol I presence indicate that nucleolin acts as an early effector in the adaptive downregulation of rDNA transcription under compression.

Next, we assessed whether nucleolin translocation triggered by cell compression affects rRNA processing. Three of the four rRNAs—18S, 5.8S, and 28S—are produced from a single long polycistronic transcript, the 47S, synthesized by RNA polymerase I (see Figure 4J) at the interface between the fibrillar center (FC) and the dense fibrillar component (DFC), implying extensive processing (cleavage) to generate the mature rRNAs. Nucleolin localizes mainly to the DFC and its peripheral region[10], where many of these key RNA cleavage steps take place.To determine whether nucleolin displacement under compression affects pre-rRNA processing, we examined pre-rRNA processing intermediates by northern blotting. Northern blot analysis revealed perfectly reproducible defects (N=4) in pre-rRNA processing under compression, including reduced levels of the primary 47S transcript and accumulation of 47S/45S precursors (Figure 4E,F,I), consistent with impaired rDNA transcription and early processing. The most pronounced changes were the accumulation of 21S and 21S-C precursors together with a reduction in 18S-E, indicating inhibition at processing sites C and E (Figure 4F,I,J). Accumulation of the 30S intermediate further suggested inhibition at site A0. As these steps contribute to small-subunit (SSU) maturation, the data indicate a preferential disruption of the SSU processing pathway upon compression. In contrast, only modest changes were observed along the large-subunit pathway, including accumulation of 32S and reduction of 12S (Figure 4G,I). Mature 18S and 28S rRNA levels remained unchanged (Figure 4H), consistent with the long half-lives of mature ribosomes. Together, these findings link nucleolin translocation under mechanical compression to both reduced rDNA transcription and impaired early pre-rRNA processing, revealing coordinated regulation of ribosome biogenesis.

### Nucleolin redistribution occurs independently of classical post-translational regulation

Nucleolin localization has been reported to be regulated by multiple post-translational modifications (PTMs), including phosphorylation, SUMOylation, methylation, glycosylation, and ADP-ribosylation [35,36]. We first focused on phosphorylation changes as these were observed to be associated with nucleolin translocation out of nucleoli [37]. Because these biochemical assays require larger cell numbers than accessible with the mechanical compression systems, we used hyperosmotic shock as an orthogonal perturbation that recapitulates nucleolin translocation. We observed that exposure to 450 mOsm hyperosmotic shock induced rapid nucleolin displacement from the nucleolus to the nucleoplasm within seconds, accompanied by nucleoplasmic foci formation and an ∼40% reduction in nucleolar nucleolin intensity (Figure 5A,B). Moreover, the internal ring-like patterned organization was also diminished, reproducing the phenotype observed under mechanical compression (Figure 5B). To determine whether nucleolin phosphorylation changes during this translocation, RFP-nucleolin was immunoprecipitated using RFP-Trap magnetic beads (Figure 5C) and analyzed using Pro-Q Diamond phosphoprotein staining together with SYPRO Ruby total protein staining for normalization. Across three independent biological replicates, no detectable changes in nucleolin phosphorylation were observed following hyperosmotic shock (Figure 5D).

**Figure 5.**
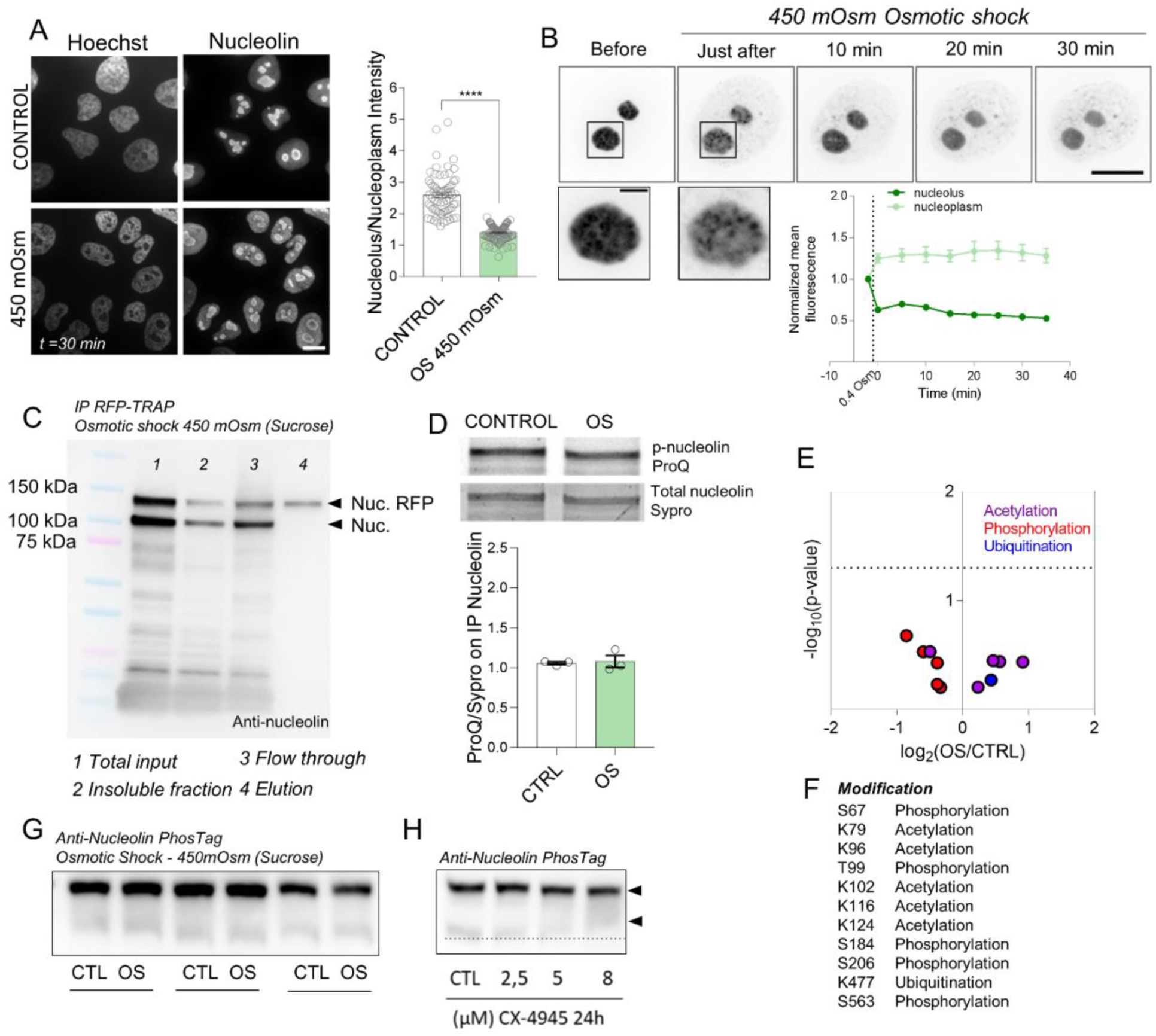
Nucleolin redistribution occurs independently of classical post-translational regulation. A) Representative images of nucleolin staining and Hoechst showing translocation of nucleolin to the nucleoplasm under hyperosmotic shock (450 mOsm sucrose). Quantification depicts the change in nucleolus-to-nucleoplasm intensity ratio. (Scale bar=10 µm, N=3 independent experiments with n>20 per condition per experiment; **** p<0.0001 in Mann-Whitney test) B) Time series of inverted fluorescence images of RFP-nucleolin–expressing cells under hyperosmotic shock, showing nucleolin accumulation in the nucleoplasm, foci formation, and reduced nucleolar intensity. Quantification over time confirms reciprocal intensity changes in the nucleolus and nucleoplasm. (Scale bar=10 µm and 2µm for zoom image, N=3 independent experiments with n=7 per condition per experiment. Each time point is represented by mean+SEM) C) Immunoprecipitation blot performed with RFP-TRAP on RFP-nucleolin cells exposed to osmotic shock. D) Purified RFP-nucleolin samples from CTL and OS conditions were resolved by SDS–PAGE and stained with Pro-Q and SYPRO, showing no detectable difference in overall nucleolin phosphorylation levels. (N=3 independent experiments). E) PTM proteomic analysis of RFP-nucleolin shows no significant differences between CTL and OS conditions for the analysed sites (log₂ fold change vs-log_10_ (p-value)). F) Representative list of nucleolin PTMs sites identified across three replicates by mass spectrometry. G) Anti-nucleolin staining visualized on Phos-tag gel on 3 independent experiments of CTL vs OS (450mOsm hyperosmotic shock) shows no clear shift of bands between CTL and OS. H) CK2 inhibition with CX-4945 induces a mild shift in nucleolin migration, indicated by the band shifts (arrows) visualized by anti-nucleolin staining on Phos-tag gels.

To probe the phosphorylation status of NCL with site-specific resolution, we next performed mass spectrometry–based quantitative phosphoproteomics. Several previously reported nucleolin phosphosites were detected (Figure 5E,F), but differential analysis revealed no statistically significant differences between control and osmotic shock conditions. Analysis of additional PTMs, including, acetylation, and ubiquitination, likewise revealed no detectable changes (Figure 5E,F; Figure S2B).

To ensure that the observations obtained under hyperosmotic shock were not specific to this perturbation, we analyzed nucleolin phosphorylation under mechanical compression using Phos-tag SDS–PAGE. Phos-tag analysis performed under the same compression modalities used above—axial confinement and in-plane compression—revealed no significant changes in nucleolin mobility compared with controls (Figure 5G, Figure S2A). Inhibition of CK2, a kinase known to phosphorylate nucleolin and regulate its subnuclear distribution [38,39], with CX-4945 produced a modest shift in nucleolin migration, confirming assay sensitivity (Figure 5H). Consistent with these findings, CK2 inhibition did not significantly alter the fraction of cells exhibiting nucleoplasmic nucleolin foci during compression (Figure S2C). All together, these results suggest that nucleolin translocation during mechanical perturbation occurs independently of detectable PTM changes, pointing to alternative regulatory mechanisms.

### Nuclear volume loss emerges as a biophysical regulator of the nucleolar response

As a biomolecular condensate, the nucleolus is sensitive to its local physicochemical environment and exhibits differential stability in response to biophysical perturbations [23,24]. Because both axial confinement and hyperosmotic shock produced an identical nucleolin translocation phenotype, we considered the factor they share in common: acute loss of nuclear volume. We therefore hypothesized that rapid reductions in nuclear volume trigger the observed nucleolar response. Previous studies indicated that gradual mechanical loading allows cells to adapt to changes in membrane tension and activate osmotic compensation mechanisms, thereby minimizing abrupt volume loss [40,41]. To isolate the contribution of volume reduction to nucleolin dynamics, we applied progressive axial confinement, gradually increasing strain over a period of 1 hour (Figure 6A).

**Figure 6.**
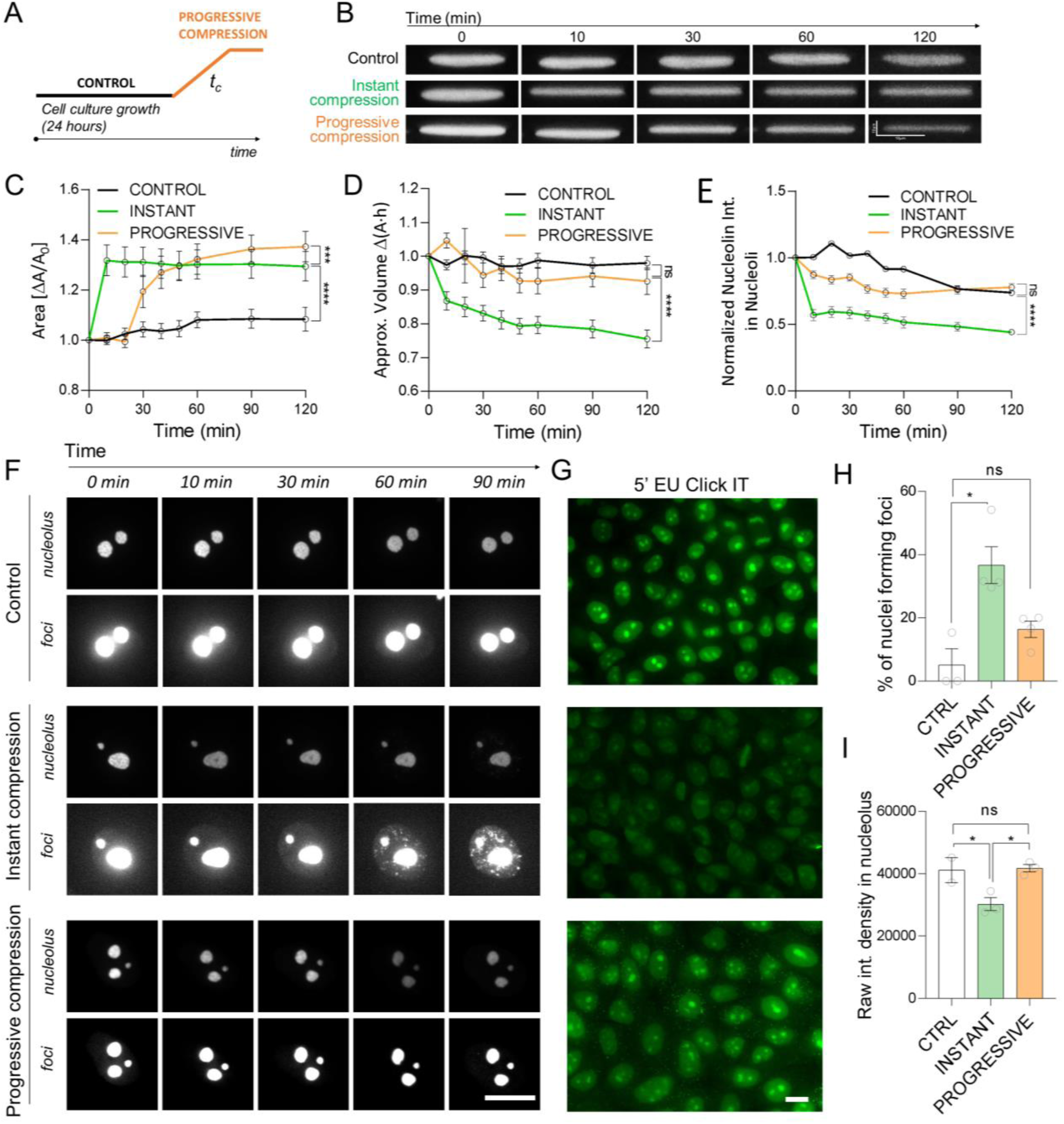
Nuclear volume loss emerges as a biophysical regulator of the nucleolar mechano-response. A) Schematic representation of the strain rate of deformation during progressive compression. B) Representative time-lapse confocal images (XZ) of MCF10A cells expressing Geminin-GFP, used to quantify nuclear morphology changes during instant and progressive compression. Time t=0 corresponds to the last pre-compression frame; compression was applied at t∼5 min, with the first post-compression image acquired at t=10 min (images acquired every 10 min). (Scale bar=10µm). C) Time-lapse quantification of nuclear area reveals a sudden deformation upon instant compression, and a gradual increase in nuclear area during progressive compression. (N=3 independent experiments, with n>3 per condition per experiment; **** p<0.0001 and *** p<0.001 in one-way ANOVA test. Each time point is represented by mean+SEM.) D) Quantification of approximate nuclear volume shows a loss under instant compression, whereas progressive compression results in no significant volume change. (N=3 independent experiments, with n>3 per condition per experiment; **** p<0.0001 in one-way ANOVA test. Each time point is represented by mean+SEM.) E) Quantification of nucleolin intensity change in nucleoli over time shows that progressive compression has minor reduction as compared to significant decrease under instant compression. (N>3 independent experiments, with n>3 per condition per experiment; **** p<0.0001 in one-way ANOVA test. Each time point is represented by mean+SEM.) F) Representative time series images of RFP-nucleolin cells exposed to instant vs progressive compression showing that progressive strain rate limits appearance of speckles indicating minimal nucleolin translocation to nucleoplasm under compression. Contrast adjusted images of the same nucleoli were depicted in each condition to visualize nucleolar morphology and formation of nucleolin foci in the nucleoplasm. (Scale bar=10 µm). G) Representative images of 5-EU Click iT assay showing decrease of nascent rRNA levels during instant compression and no significant changes for progressive compression. (Scale bar=10 µm) H) Quantification of the percentage of nuclei displaying nucleoplasmic speckles shows no significant difference under progressive compression compared to control. (N=3 independent experiments, with n>11 per condition per experiment; * p<0.05 in one-way ANOVA test) I) Quantification of 5-EU signal intensity within the nucleolus further confirms decrease in nascent rRNA levels during instant compression and no significant changes for progressive compression. (N=3 independent experiments with n>40 analyzed per condition per experiment; *p<0.01 in one-way ANOVA test)

MCF10A FUCCI cells were used to visualize distinct nuclear deformation patterns, using their diffuse, non–chromatin-bound fluorescence, which is well suited for 3D live imaging (Figure 6B). Instant compression of cells triggered an abrupt increase in projected nuclear area and a reduction in nuclear volume, whereas progressive compression induced gradual deformation of the nuclear area with minimal volume loss (Figure 6C, 6D, Figure S3A). Strikingly, progressive compression significantly reduced nucleolin depletion from the nucleolus upon compression (Figure 6E,F). Additionally, only ∼18% of nuclei formed fluorescent nucleolin foci in the nucleoplasm under progressive compression (not significant against control), compared to ∼38% observed under instant compression (Figure 6F, 6H). Consistent with the absence of significant changes in nucleolar nucleolin abundance, EU–click-IT labelling confirmed that nascent rRNA levels within the nucleolus during progressive compression were unchanged relative to controls (Figure 6G), indicating that the rate of rRNA production remains unperturbed under progressive compression and without displacement of nucleolin. Together, these results indicate that gradual nuclear deformation prevents nucleolin translocation and preserves nucleolar transcription.

To further test the hypothesis of a volume-dependent mechanism, we subjected cells to graded hyperosmotic shock (300, 450, 600 mOsm sucrose). Nucleolin translocation increased proportionally with hyperosmolarity (Figure S3B). This dose-dependent response reinforces the hypothesis that nuclear volume loss strongly influences nucleolin translocation. In all previous experiments, cells were analyzed in an unsynchronized state. To test whether increasing nuclear size modulates this response, we performed instant compression on monolayers of cells co-expressing geminin–GFP and RFP-nucleolin, synchronized to the S–G2 phase of the cell cycle, characterized by increased geminin expression (Figure S3C). Cells in S–G2 phase possess larger nuclei and therefore experience stronger strain and deformation upon mechanical confinement compared to unsynchronized monolayers. Consistently, compression of synchronized monolayers resulted in a stronger nucleolin response, reflected by a higher percentage of cells forming nucleolin foci in the nucleoplasm (Figure S3D). Altogether, progressive compression along with the hyperosmotic shock experiments, demonstrated that nucleolin redistribution under mechanical compression depends on the rate of nuclear deformation and the associated loss of nuclear volume, highlighting the sensitivity of the nucleolus to rapid biophysical perturbations of its environment.

### Compression-induced nucleolar adaptation involves localized H4K20me3 remodeling but not H3K9-dependent silencing

Mechanical compression rapidly reduced rRNA synthesis on the minute time scale, coinciding with nuclear volume loss and nucleolin translocation from the nucleolus. We next examined whether longer-term nucleolar adaptation involves epigenetic remodeling of rDNA chromatin, focusing on established silencing mechanisms characterized by enrichment of repressive histone marks such as H3K9me2/3 [42] and H4K20me3 [43,44] at rDNA loci. First, we used expansion microscopy to examine chromatin organization under mechanical compression. Expansion microscopy revealed pronounced chromatin reorganization visualized with Hoechst staining (Figure S4A). Globally, chromatin appeared more decondensed, with euchromatin expanding into larger nuclear domains. At the same time, compressed cells displayed discrete heterochromatin inclusions associated with nucleoli (Figure S4B). Quantification revealed a marked increase in the proportion of nuclei containing nucleolar heterochromatin patches compared with uncompressed controls, where ∼50% of nuclei lacked such domains (Figure S4C).

Next, we examined whether mechanical compression alters the distribution of repressive histone marks associated with rDNA silencing. Although compressed cells occasionally displayed H3K9me3-enriched heterochromatin rings surrounding nucleoli, quantitative analysis revealed no enrichment of H3K9me3 at the nucleolar periphery and instead showed an overall reduction in nuclear H3K9me3 levels after 2 h of compression (Figure S4E). Similarly, the related repressive mark H3K9me2 was significantly downregulated following compression (Figure S4D) as previously observed for compressed tissues [45]. We next asked whether the NoRC component BAZ2A/TIP5 becomes enriched at nucleoli upon compression. No increase in nucleolar-associated BAZ2A signal was detected (Figure S4F), in contrast to H3K9-dependent rDNA silencing conditions that involve BAZ2A recruitment to rDNA [34]. Notably, quantification of the nucleolar-associated BAZ2A signal revealed a modest decrease at early time points (30 min), with no detectable difference at later time points (2 hrs; Figure S4F) compared to control. Overall, these results suggest that compression-induced rDNA repression does not primarily rely on the canonical H3K9–NoRC silencing pathway.

An alternative rDNA silencing pathway involves H4K20me3 deposition mediated by recruitment of the histone methyltransferase SUV420H2 to nucleoli by the long non-coding RNA PAPAs, as previously described during growth arrest and osmotic stress [43,44]. Immunofluorescence analysis demonstrated a global reduction in nuclear H4K20me3 levels upon 2h of mechanical compression (Figure 7A). Despite this decrease, SUV420H2 displayed significant enrichment within nucleoli in compressed cells (Figure 7B). Notably, SUV420H2 localization was not altered at earlier time points (30 min post-compression; Figure S4G), indicating that its nucleolar accumulation occurs as a delayed response to sustained mechanical stress. To assess whether the lncRNA PAPAs mediates SUV420H2 enrichment in the nucleolus under mechanical stress, we quantified the abundance of PAPAs transcript by RT-qPCR. PAPAs transcript abundance increased after 2 hrs of compression and declined 1 hr after stress reversal (Figure 7C). Together, these results indicate that activation of the PAPAs–SUV420H2 pathway occurs at later stages of the mechanical response and represents a potential mechanism contributing to compression-induced rDNA repression.

**Figure 7.**
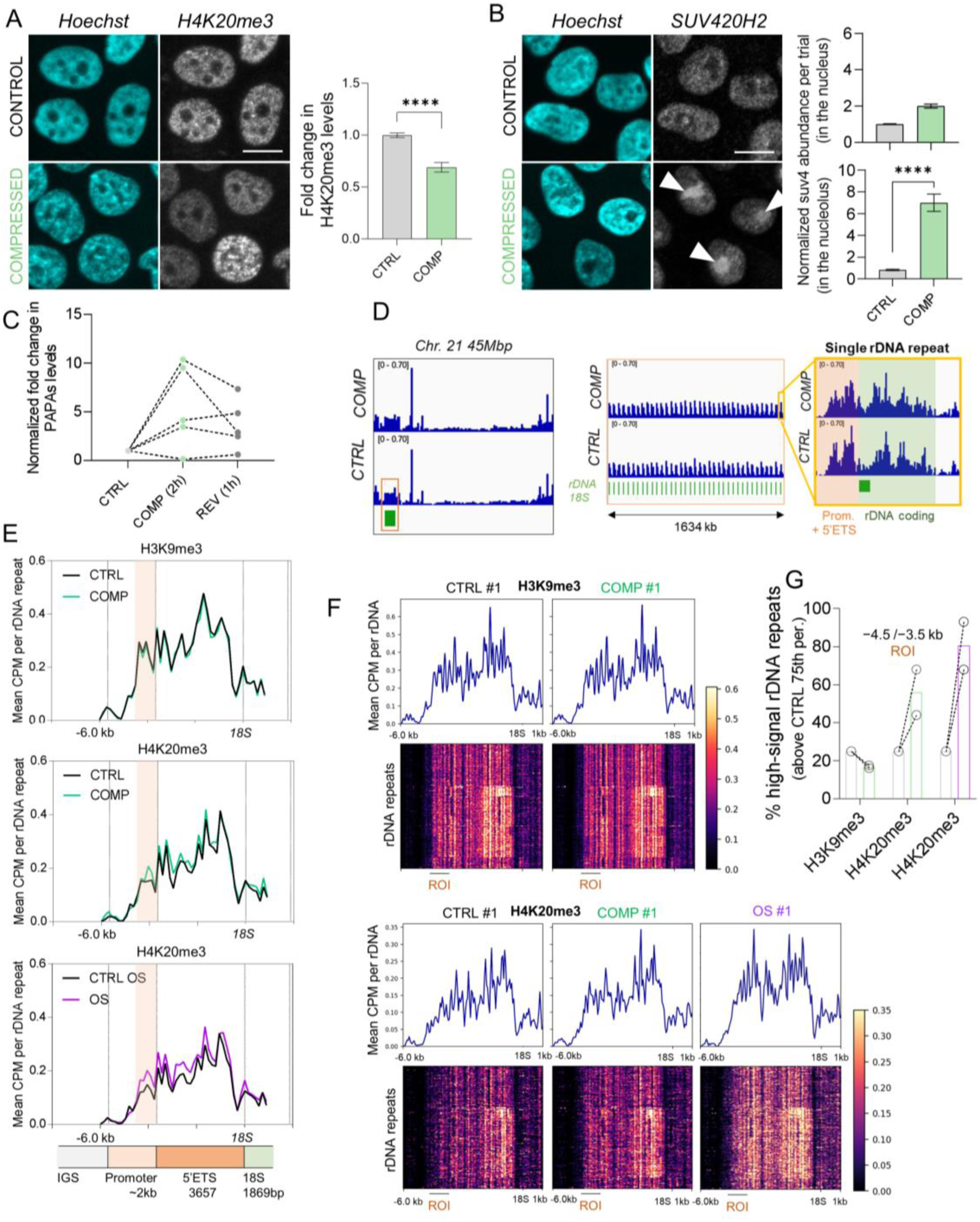
Compression induced nucleolin translocation is coupled with global loss of heterochromatin and nucleolar enrichment of SUV420H2 methyltransferase. A) Representative images of staining with anti-H4K20me3 and Hoechst in control and compressed samples. Quantification in whole nuclei showed decrease in H4K20me3 levels under compression. (Scale bar=10 µm, N=3 independent experiments with n>35 cells per condition per experiment; ****p<0.0001 in unpaired t test with Welch’s correction. B) Immunofluorescence staining and quantification of SUV420H2 intensity shows no significant changes in the nucleus and enrichment in the nucleolus (arrows) under compression. (Scale bar=10µm, N=3 independent experiments with n>25 nuclei and n>40 nuceloli per condition per experiment; ****p<0.0001 in Mann-Whitney test) C) RT qPCR results depicting the total cellular levels of lncRNA PAPAs in control and compressed conditions (N=5 independent experiments). D) IGV tracks showing CUT&Tag signal across a representative rDNA array on chromosome 21. The repetitive organization of rDNA units is visible in the middle panel, with a zoom-in highlighting signal distribution across a single repeat (right). The promoter with 5’ETS and coding regions are indicated, illustrating signal enrichment within the rDNA unit. E) Mean Cut&Tag CPM profiles for H3K9me3 and H4K20me3 are shown from −6 kb to the 18S start site, representing the mean of two biological replicates. The highlighted region denotes the region of interest (ROI; −4.5 to −3.5 kb), where the strongest changes upon compression and osmotic shock are observed. Vertical dotted lines correspond to the rDNA organization scheme below and indicate the promoter, 5′ETS, and 18S start site. F) Repeat-resolved chromatin profiles across rDNA under different conditions. Heatmaps and corresponding mean CPM profiles for H3K9me3 and H4K20me3 across individual rDNA repeats are shown for representative single replicates under control (CTRL), compression (COMP), and osmotic shock (OS) conditions. Signal is plotted from −6 kb upstream to the 18S start site. Each row in the heatmap represents an individual rDNA repeat, highlighting repeat-level heterogeneity in chromatin enrichment. The region of interest (ROI; −4.5 to −3.5 kb) is indicated and corresponds to the promoter region where the most pronounced changes are observed. G) Percentage of rDNA repeats with signal exceeding a CTRL-defined threshold (75th percentile) within the ROI (−4.5 to −3.5 kb) is shown for each condition. Points represent individual biological replicates, and bars indicate mean values. This approach captures shifts in the fraction of high-signal repeats across conditions.

To determine whether mechanical compression induces locus-specific epigenetic changes at rDNA, we performed CUT&Tag profiling for H3K9me3 and H4K20me3. To enable quantitative analysis of chromatin organization across ribosomal DNA repeats, we mapped CUT&Tag reads to the telomere-to-telomere human reference genome (T2T-CHM13) [46], which resolves previously unmapped repetitive rDNA regions (Figure 7D). CUT&Tag signal tracks were normalized to counts per million (CPM) to enable comparison between conditions. Given that transcription initiates at the rDNA promoter/5′ETS and that nucleolin is enriched at this region under basal conditions but rapidly dissociates upon compression [34], we focused our analysis on this promoter-proximal domain to assess chromatin organization at the site of transcription initiation. Pairwise comparison revealed similar spatial enrichment profiles across biological replicates within the rDNA promoter/5′ETS region, despite inter-replicate differences in global signal amplitude (Figure S5A). In contrast to the global chromatin decondensation observed by immunofluorescence, CUT&Tag revealed highly similar H3K9me3 enrichment profiles across the rDNA promoter region, with control and compressed samples displaying nearly indistinguishable signal distributions when averaged across rDNA repeats and independent biological replicates (Figure 7E,F; Figure S5B). These observations indicate that compression-induced chromatin remodeling at rDNA occurs independently of major H3K9me3 redistribution..

Consistent with the activation of the PAPAs–SUV420H2 pathway, we next examined H4K20me3 distribution across rDNA. H4K20me3 showed subtle, reproducible but spatially coherent remodeling within the upstream rDNA promoter region (∼−4.5 kb to −3.5 kb relative to the 18S start site), with modest signal alterations detected at the promoter and within the 5′ external transcribed spacer (5′ETS) - regions linked to transcriptional regulation of rDNA (Figure 7E, F ; Figure S5B). Consistently, changes at the rDNA promoter were also observed following osmotic shock, used here as a positive control for activation of this previously described regulatory pathway (Figure 7E, F; Figure S5B). To quantify these changes at the repeat level, we calculated the fraction of rDNA repeats exhibiting signal above a threshold defined by the 75th percentile of the CTRL distribution within the ROI (Figure 7G). This analysis revealed only minor changes (∼5%) in the proportion of high-signal repeats for H3K9me3 upon compression, consistent with the absence of coordinated enrichment. In contrast, H4K20me3 displayed a marked increase (∼25%) in high-signal repeats under compression, which was further enhanced under osmotic shock (∼55%), a condition known to more strongly repress rDNA transcription. Together, these findings define a temporally distinct, second mode of nucleolar adaptation in which PAPAs–SUV420H2–dependent epigenetic remodeling selectively targets a narrow rDNA promoter region. This delayed response (∼2 hrs) follows the rapid nucleolin-mediated adaptation occurring within minutes of compression and is characterized by consistent enrichment of H4K20me3 across rDNA repeats under both compression and osmotic stress.

## DISCUSSION

Mechanobiology has emerged as a central regulator of cellular behavior, with mechanical forces known to influence processes ranging from cytoskeletal organization and membrane signaling to nuclear architecture and gene expression. An increasing number of intracellular structures and organelles have been shown to sense and respond to mechanical cues [47], yet whether biomolecular condensates participate directly in mechanosensitive regulation remains largely unexplored. Here, we identify the nucleolus—the primary site of ribosome biogenesis—as a mechanosensitive organelle, placing it among a growing class of intracellular compartments whose structure and function are directly modulated by mechanical forces. Mechanical compression induces reversible reduction in rDNA transcription, revealing a previously unrecognized pathway linking mechanical forces to the regulation of ribosome production. Such compressive stresses commonly arise in physiological contexts including epithelial crowding, tissue morphogenesis, cell migration through confined environments [48,49], and in pathological states such as tumor growth and metastasis, where cells frequently experience increased mechanical pressure and transient reductions in cellular and nuclear volume [50]. The rapid and reversible nucleolar response described here therefore suggests that mechanical regulation of nucleolar activity may contribute to the dynamic adaptation of ribosome biogenesis, and more broadly biosynthetic activity[51,52], to such volume-dependent physical constraints in tissues.

Unlike classical nucleolar stress responses that lead to nucleolar disruption and activation of p53 signaling, nucleolin redistribution occurs rapidly upon nuclear volume reduction and precedes detectable p53 activation, indicating that it represents an early mechanosensitive response rather than a downstream stress pathway. Therefore, our findings identify nucleolin as a key mechano-responsive regulator of ribosome biogenesis whose nucleolar localization rapidly and reversibly adapts to mechanical compression. Nucleolin spans multiple nucleolar subcompartments—including the DFC (with its peripheral regions) [9], and extends towards the GC [53] — where it supports successive steps of ribosome biogenesis [54,55]. Consistent with its multifunctional role, nucleolin translocation under mechanical compression coincided with reduced rDNA transcription and detectable changes in early pre-rRNA processing, indicating that nucleolin redistribution impacts ribosome biogenesis across multiple levels of the pathway. Notably, we observe modulation of processing events associated with small ribosomal subunit formation, which occurs predominantly in the nucleolin-enriched peripheral dense fibrillar component (pDFC) [10], revealing a previously unrecognized link between mechanical stress and rRNA processing. Together, these findings suggest that mechanical redistribution of nucleolin functionally resembles partial nucleolin depletion, which has been shown to rapidly reduce rDNA transcription and modulate ribosome biogenesis [33,34]. These observations position nucleolin redistribution as an early mechanosensitive event that precedes subsequent epigenetic repression of rDNA, consistent with the PAPAS–SUV420H2 pathway that deposits H4K20me3 on rDNA during growth-constrained states such as highly dense cultures [44], a condition that mechanically parallels the confinement imposed by in-plane compression in our experiments. This temporal separation suggests that mechanical perturbations first trigger rapid, reversible changes in nucleolar organization through nucleolin redistribution, and are subsequently stabilized by promoter-specific epigenetic modifications that reinforce longer-term repression of rDNA transcription. Importantly, this redistribution occurs without detectable large-scale changes in nucleolin post-translational modifications but is highly sensitive to rapid reductions in nuclear volume, suggesting that nucleolin localization responds primarily to the physicochemical consequences of compression rather than to canonical signaling pathways.

As a biomolecular condensate whose assembly depends on concentration-dependent multivalent interactions [56,57], the nucleolus is inherently sensitive to changes in its physicochemical environment, including macromolecular concentration, ionic conditions, and molecular crowding. Mechanical compression and the associated reduction in nuclear volume are expected to rapidly increase the effective concentration of nucleoplasmic components, thereby altering the phase behavior and internal organization of nucleolar assemblies. Such changes can influence the partitioning of proteins between nucleolar subcompartments and the surrounding nucleoplasm without requiring active biochemical signaling. In this framework, the observed redistribution of nucleolin may reflect a shift in its phase partitioning within the nucleolar network in response to compression-induced changes in concentration and crowding. In a multicomponent condensate such as the nucleolus, changes in global concentration are not expected to uniformly promote phase separation of all components. Instead, differential interaction strengths and competition between binding partners will lead to selective partitioning or even exclusion of specific proteins.

An important question, however, is why nucleolin appears particularly sensitive to these perturbations whereas many other nucleolar proteins remain stably associated with the nucleolus. Insights from minimal in vitro reconstitution systems suggest that different nucleolar proteins exhibit distinct saturation concentrations and phase transition thresholds [58], determined by intrinsic molecular features such as the number and distribution of intrinsically disordered regions, multivalent RNA-binding domains, charge patterning, and interaction valency with rRNA and other scaffold proteins. As a result, global physicochemical changes affecting the nucleolus—such as those produced by nuclear volume reduction—are expected to differentially alter the partitioning behavior of its components, causing some proteins to exit condensates more readily than others. Redistribution of nucleolin may also involve changes in its interaction balance between the nucleolus and other nuclear compartments, such as chromatin. Future work will be required to determine which physicochemical parameters associated with nuclear compression—such as macromolecular crowding, ionic composition, or RNA availability or changes in interaction landscapes across nuclear compartments —most directly control nucleolar phase behavior and the selective redistribution of nucleolin. From a broader perspective, because mechanical forces frequently reshape cellular and nuclear volumes, similar principles may extend to many biomolecular condensates throughout the cell, suggesting that mechanical regulation of intracellular volume may represent a general physical mechanism linking mechanical forces to condensate organization and function.

## AUTHOR CONTRIBUTIONS

YS, KE, SB, LP, A-SR, PN, LB, MED: Performed all experiments and analyses except if otherwise stated.

LB, YC: MS based proteomics (experiment and analysis; figure preparation)

FE: Half-FRAP (analysis and theoretical model, figure preparation)

EM and YS: performed Cut and Tag experiments

LW and CZ: RNA biochemistry; pre-rRNA analysis

DLJL: RNA biochemistry and processing (funding acquisition; figure preparation)

M.E.D: conceived, acquired funding, supervised the study, and wrote the paper.

All authors edited the manuscript.,

## ACKNOWLEDGEMENTS

We thank Alexei Grichine, Mylène Pezet, Jacques Mazzega, Solenne Dufour, and Florence Appaix from the IAB microscopy platform for assistance with imaging and FACS experiments. We thank Bastien Touqet (Tardieux lab) and Karin Sadoul for training and discussions regarding expansion microscopy. We also thank Guillermo Orsi for discussions regarding CUT&Tag experiments and analysis.

K.E. received doctoral scholarship from IRGA program from University of Grenoble Alpes. S.B. received doctoral scholarship from La Ligue Contre le Cancer foundation. Research in the team of M.E.D was supported by Agence Nationale de la Recherche (grant ANR-17-CE13-0006), Foundation ARC (ARCPJA2023090007058), CNRS IEA program (Gradient), and Espoir Isere foundation.

High-throughput sequencing was performed by the ICGex NGS platform of the Institut Curie supported by the grants ANR-10-EQPX-03 (Equipex) and ANR-10-INBS-09-08 (France Génomique Consortium) from the Agence Nationale de la Recherche (“Investissements d’Avenir” program), by the ITMO-Cancer Aviesan (Plan Cancer III) and by the SiRIC-Curie program (SiRIC Grant INCa-DGOS-465 and INCa-DGOS-Inserm_12554). Data management, quality control and primary analysis were performed by the Bioinformatics platform of the Institut Curie.

Research in the laboratory of D.L.J.L. was supported by the Belgian Fonds de la Recherche Scientifique (F.R.S.–FNRS), EOS (CD-INFLADIS, 40007512), Région Wallonne (SPW EER) Win4SpinOff (RIBOGENESIS), the COST Action TRANSLACORE (CA21154), the European Joint Programme on Rare Diseases (EJP-RD): RiboEurope and DBAGeneCure, and the Marie Skłodowska-Curie Actions Doctoral Network (MSCA-DN) EURECA (https://eureca-dn.com/)

The proteomic experiments were partially supported by Agence Nationale de la Recherche under projects ProFI (Proteomics French Infrastructure, ANR-10-INBS-08 & ANR-24-INBS-0015) and GRAL, a program from the Chemistry Biology Health (CBH) Graduate School of University Grenoble Alpes (ANR-17-EURE-0003).

## Notes

### Competing Interest Statement

The authors have declared no competing interest.

