## Supplementary material for "The nucleolus is a mechanosensitive condensate that adapts ribosome biogenesis to mechanical forces": Supp Info

Figure S1

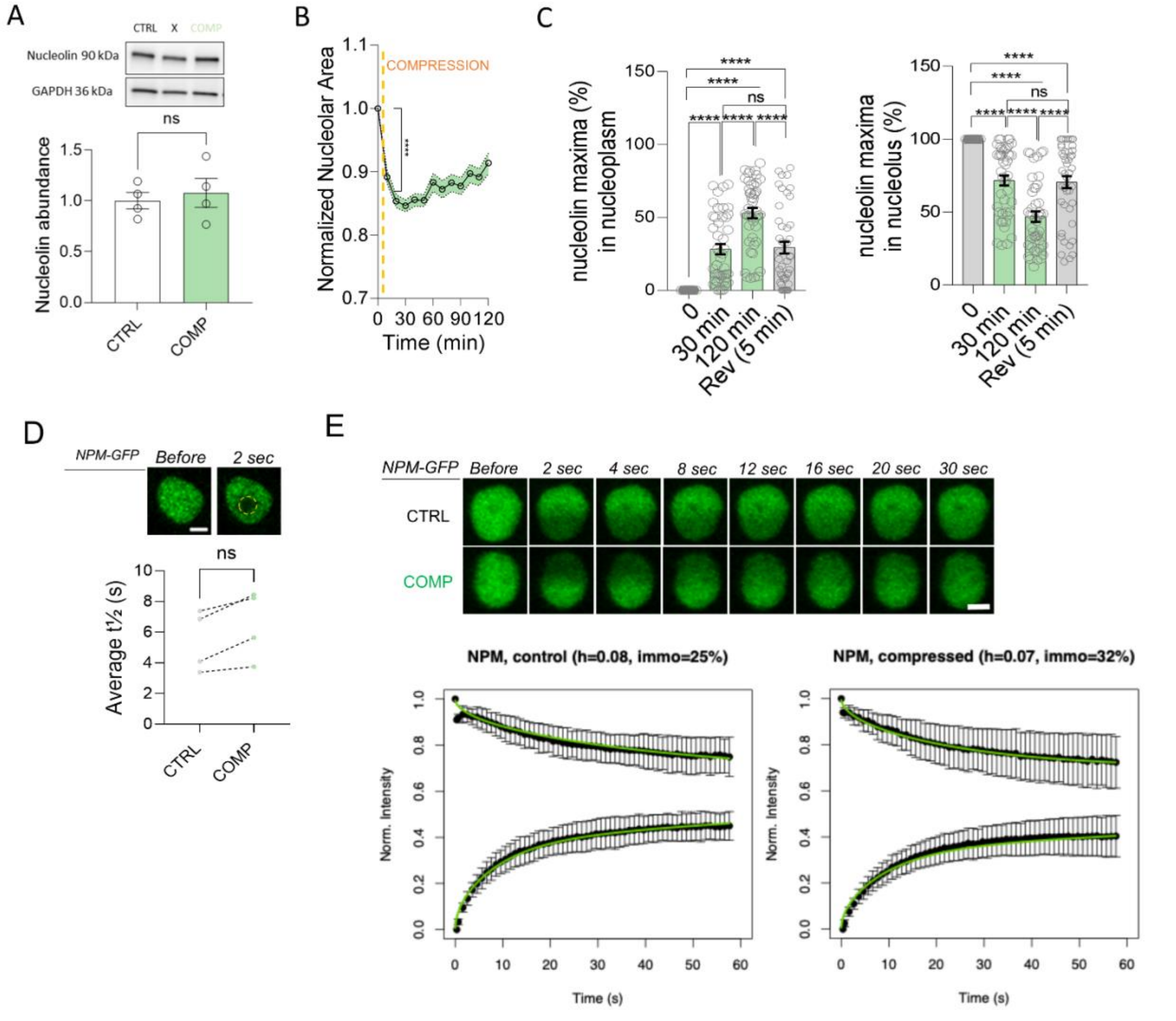


***Figure S1 in relation to Figure 3***

*A) Total abundance of nucleolin does not change under axial compression as shown by WB quantification (N=4 of independent experiments; p=n.s. in unpaired t test with Welch's correction).*

*B) Normalized changes in nucleolar area upon compression show a 10% deformation within 10 minutes of compression and a partial recovery during the following 110 minutes (pillar height= 4µm). (N=3 independent experiments, with n=40 nucleoli analyzed per condition per experiment; **** p<0.0001 in Mann Whitney test. Each time point is represented by mean + SEM.)*

*C) Percentage of nucleolin maxima found in nucleoplasm (left panel) vs nucleolus (right panel) at the given time points before, during compression and upon release of compression. (N=3 independent experiments with n=15 nuclei analyzed per condition per experiment; **** p<0.0001 in Friedman test).*

*D) Representative NPM-GFP images acquired immediately before FRAP and 2 s after bleaching, with the bleached area marked in yellow. Quantification of the average half-time of recovery ($\tau_{1}/_{2}$) reveals no change in NPM mobility within the nucleolus under axial compression. (Scale bar=1µm, N=4 independent experiments with n>5 nucleoli analyzed per condition per experiment; p=n.s. in unpaired t test with Welch's correction).*
*E) Time-series images of NPM-GFP labeled nucleoli bleached in one half of their volume. Plots of intensity changes show mean normalized intensity in the bleached and non-bleached halves revealed recovery kinetics with $\tau$ = 9 s for both control and compressed cells. Green curves represent the three-parameter fit model indicated in the results section. (Scale bar=1 µm, N = 2 independent experiments with n = 27 nucleoli per condition within three biological replicates).*

Figure S2

A

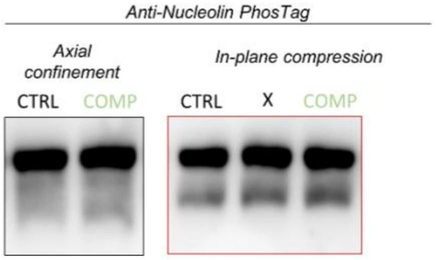


B

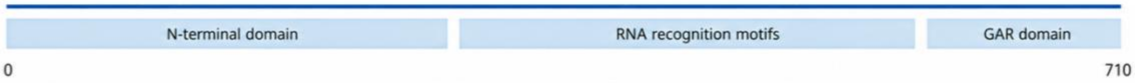


| Residue | Modification | Best psm score | Sites Confidence(%) | OS Vs Ctrl | | log2(normalized, filtered and imputed abundances) | | | | | |
|---|---|---|---|---|---|---|---|---|---|---|---|
| | | | | $log_2$(Fold Change) | p-value | Ctrl R1 | Ctrl R2 | Ctrl R3 | OS R1 | OS R2 | OS R3 |
| S67 | Phosphorylation | 57.29 | 99.8 | -0.60 | 2.95E-01 | 28.332 | 26.685 | 26.926 | 28.096 | 27.961 | 27.681 |
| K79 | Acetylation | 35 | 79.0 | 0.91 | 3.66E-01 | 17.575 | 18.185 | 18.012 | 18.006 | 14.536 | 18.491 |
| K96 | Acetylation | 33.4 | 91.2 | -0.50 | 2.95E-01 | 19.614 | 19.318 | 19.004 | 20.271 | 19.801 | 19.364 |
| T99 | Phosphorylation | 34.29 | 75.1 | -0.86 | 2.10E-01 | 20.145 | 17.882 | 19.098 | 20.440 | 19.486 | 19.767 |
| K102 | Acetylation | 31.75 | 99.9 | 0.23 | 6.24E-01 | 18.053 | 17.819 | 18.288 | 18.371 | 17.633 | 17.476 |
| K116 | Acetylation | 32.44 | 100.0 | 0.56 | 3.63E-01 | 19.446 | 19.751 | 19.875 | 19.981 | 17.977 | 19.449 |
| K124 | Acetylation | 36.34 | 77.3 | 0.46 | 3.57E-01 | 20.427 | 20.139 | 20.550 | 20.594 | 19.322 | 19.811 |
| S184 | Phosphorylation | 135.5 | 100.0 | -0.34 | 6.26E-01 | 24.683 | 26.045 | 26.296 | 24.784 | 26.697 | 26.567 |
| S206 | Phosphorylation | 148.87 | 100.0 | -0.39 | 5.86E-01 | 24.734 | 26.118 | 26.386 | 24.850 | 26.840 | 26.725 |
| K333 | Ubiquitination | 59.82 | 100.0 | N/A | | | | | | | |
| K477 | Ubiquitination | 38.89 | 93.5 | 0.43 | 5.36E-01 | 20.335 | 18.388 | 19.387 | 19.747 | 17.961 | 19.123 |
| S563 | Phosphorylation | 56.79 | 100.0 | -0.39 | 3.73E-01 | 26.645 | 26.567 | 26.865 | 26.765 | 27.142 | 27.331 |
| S619 | Phosphorylation | 41.62 | 100.0 | N/A | | | | | | | |

C

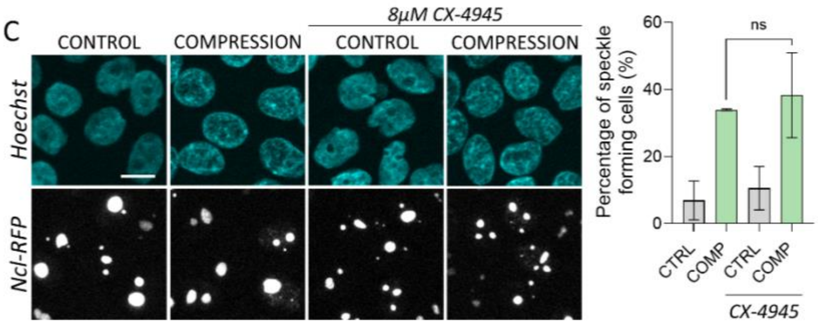


***Figure S2 in relation to Figure 5***

*A) Phos-tag staining of nucleolin demonstrates no major changes in migration pattern between control and compression condition, consistent with the absence of phosphorylation-related changes. "X" is used to mark conditions not reported in this study. (N=3 independent experiments).*

*(B) Summary of nucleolin post-translational modifications (PTMs) identified by mass spectrometry. Detected modifications included phosphorylation, acetylation, oxidation, and diglycine (GG) remnants indicative of ubiquitination. Only PTM sites with localization score >0.75*

*and reproducibly detected across replicates are shown. Quantitative comparison of PTM-containing nucleolin peptides between control and compression conditions is summarized, including site localization, log2 fold change, and statistical significance. No modified peptide displayed statistically significant abundance changes between conditions.*

*C) Representative images illustrating Nucleolin-RFP forming fluorescent foci in the nucleoplasm in both CX-4945–treated (CK2-inhibition) and untreated cells subjected to compression. Quantification of percentage of cells forming nucleolin foci depicts no significant change in nucleolin translocation. (Scale bar=10μm, N=2 independent experiments, n>94 per condition per experiment; p=n.s. in unpaired t test with Welch's correction)*

Figure S3

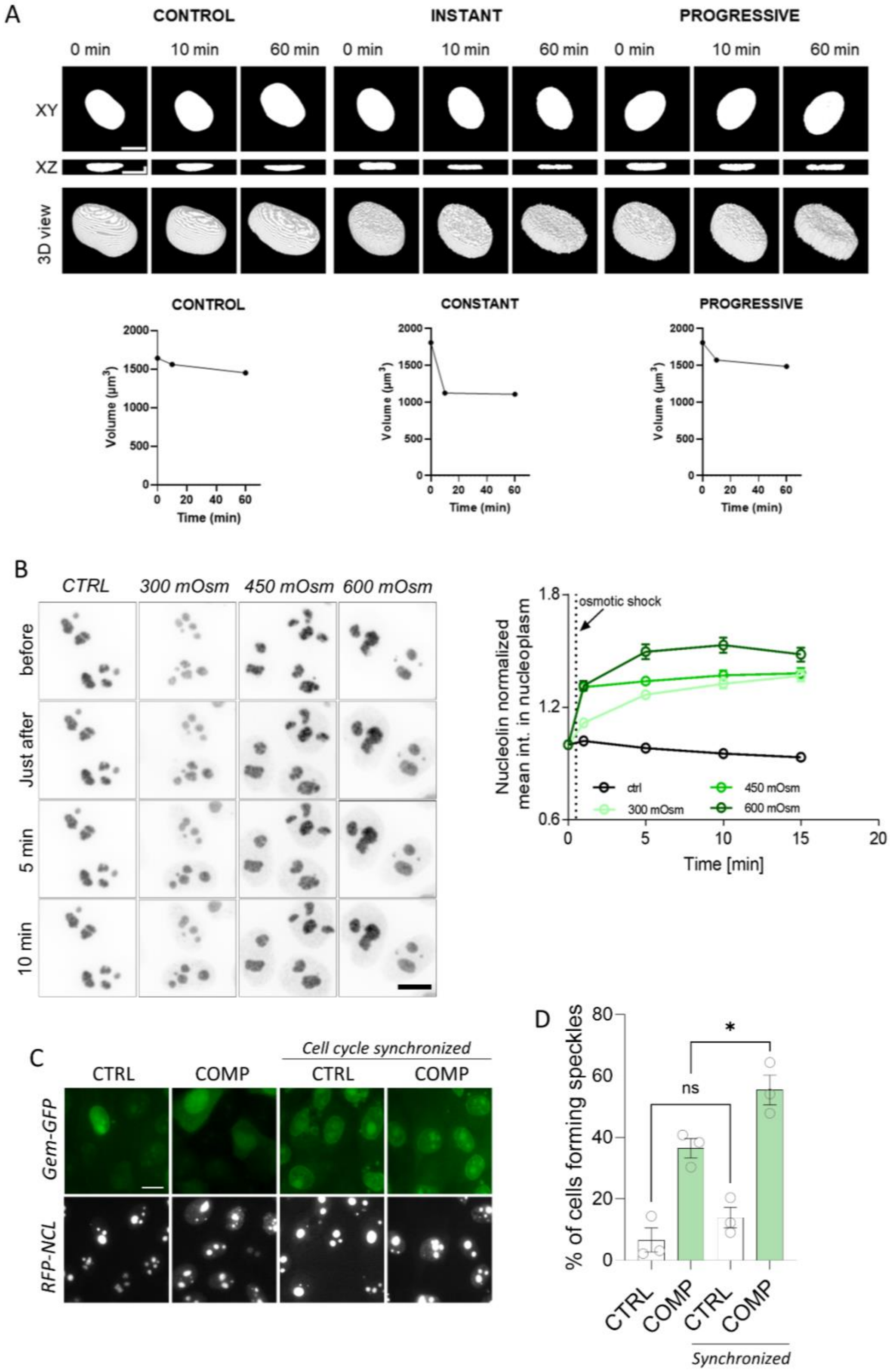

A
CONTROL
INSTANT
PROGRESSIVE
0 min
10 min
60 min
XY
XZ
3D view
CONTROL
CONSTANT
PROGRESSIVE
Volume (μm³)
Time (min)
B
CTRL
300 mOsm
450 mOsm
600 mOsm
before
Just after
5 min
10 min
osmotic shock
Nucleolin normalized mean int. in nucleoplasm
Time [min]
ctrl
300 mOsm
450 mOsm
600 mOsm
C
Cell cycle synchronized
CTRL
COMP
Gem-GFP
RFP-NCL
D
% of cells forming speckles
ns
*
Synchronized

***Figure S3 in relation to Figure 6***

*A) Representative microphotographs of XY and XZ view of the nuclei and their 3D view showing the changes in volume regulation between control, instant compression and progressive compression, and the corresponding volume change in time. Scale bar = 10 µm.*
*B) Time series of inverted RFP-nucleolin images of cells before and after exposure to modulated hyperosmotic shock (300-600mOsm). (Scale bar=10µm). Graph represents quantification showing that progressive change in nucleolin intensity in nucleoplasm in time is dependent on the magnitude of the shock. (N=2 independent experiments, n>5 nuclei per condition per experiment. Each time point is represented by mean+SEM.)*
*C) Representative images of Geminin-GFP–expressing cells synchronized to S–G2 phase with enlarged nuclei show a marked increase in nucleolin foci within the nucleoplasm. (Scale bar=10µm)*
*D) Quantification indicates that this effect is significantly higher than in compression of non-synchronized cells. (N=3 independent experiments, n>45 per condition per experiment, * p<0.05 in unpaired t test with Welch's correction)*

Figure S4

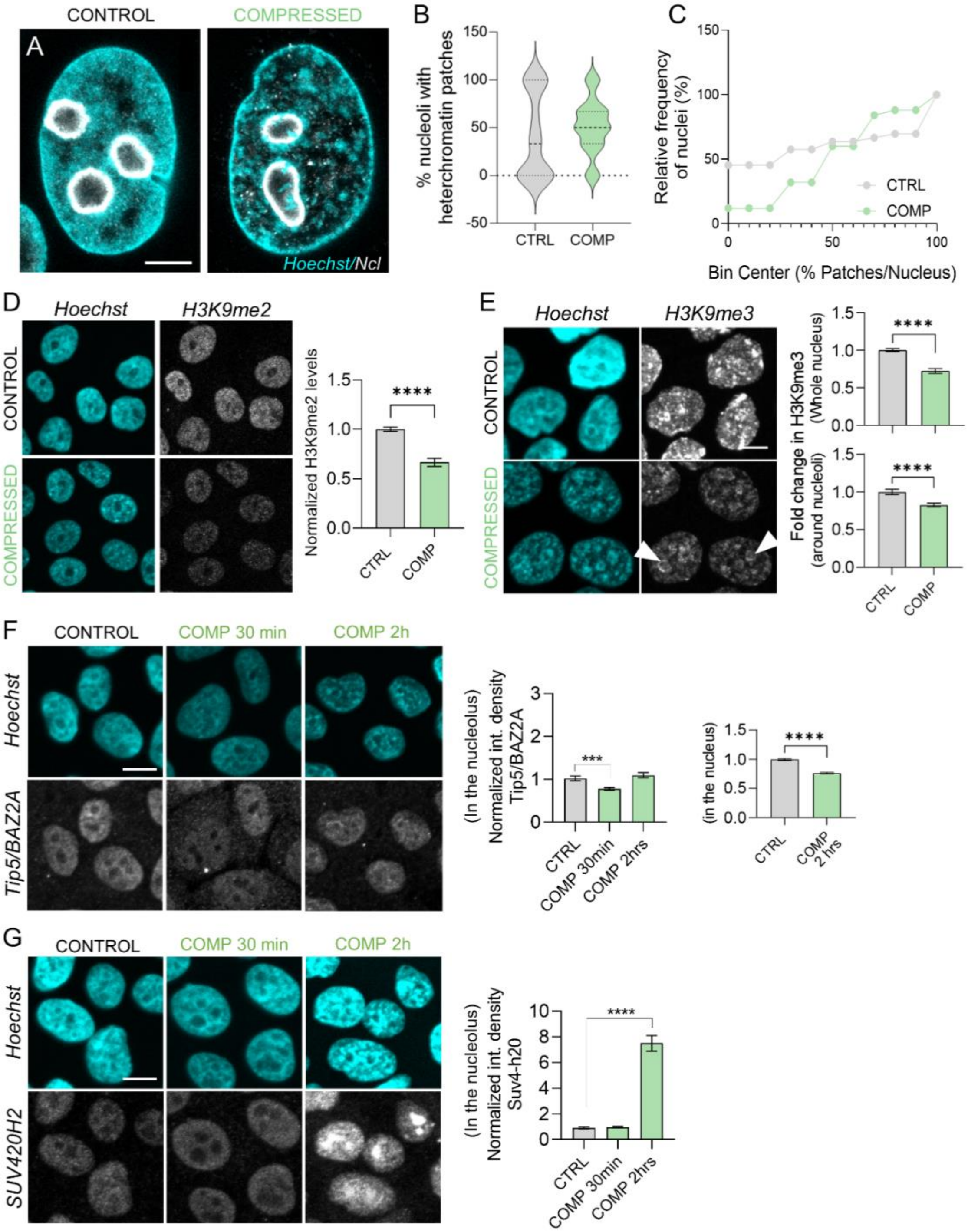

CONTROL
COMPRESSED
A
Hoechst/Ncl
B
% nucleoli with heterochromatin patches
CTRL
COMP
C
Relative frequency of nuclei (%)
Bin Center (% Patches/Nucleus)
CTRL
COMP
D
Hoechst
H3K9me2
CONTROL
COMPRESSED
Normalized H3K9me2 levels
****
CTRL
COMP
E
Hoechst
H3K9me3
CONTROL
COMPRESSED
Fold change in H3K9me3 (Whole nucleus)
Fold change in H3K9me3 (around nucleoli)
****
****
CTRL
COMP
F
CONTROL
COMP 30 min
COMP 2h
Hoechst
Tip5/BAZ2A
(In the nucleolus) Normalized int. density Tip5/BAZ2A
***
CTRL
COMP 30min
COMP 2hrs
(in the nucleus)
****
CTRL
COMP 2 hrs
G
CONTROL
COMP 30 min
COMP 2h
Hoechst
SUV420H2
(In the nucleolus) Normalized int. density Suv4-h20
****
CTRL
COMP 30min
COMP 2hrs

***Figure S4 in relation to Figure 7***

*Unless otherwise indicated, cells were analyzed after 2 hrs of compression.*

*A) Expansion microscopy images stained for nucleolin and hoechst showing presence of heterochromatin patches within the nucleolus. (Scale bar=10 μm)*

*B) Quantification of the percentage of nucleoli containing heterochromatin patches in control and compressed cells reveals a tendency of increased accumulation under compression. (N=3 independent experiments with n>5 nucleoli analyzed per condition per experiment)*

*C) Line plot showing the relative frequency distribution of nuclei exhibiting heterochromatin patches versus the percentage of patches per nucleus to reveal compression has increased distribution of heterochromatin patches per nuclei. (N=3 independent experiments with n>5 nucleoli analyzed per condition per experiment)*

*D) Representative images and quantification for H3K9me2 staining shows an overall decrease in its nuclear levels. (Scale bar=10μm, N=3 independent experiments, n>38 per condition per experiment, ****p<0.0001 in Mann-Whitney test)*

*E) Representative images of immunofluorescence staining with anti-H3K9me3 and Hoechst in control and compressed samples. Quantification of H3K9me3 intensity in whole nuclei and within a 2-pixel boundary around the nucleolus (arrows) showed an overall decrease in H3K9me3 levels under compression. (Scale bar=10μm, N=3 independent experiments with n>26 cells per condition per experiment; ****p<0.0001 in Mann-Whitney test)*

*F) Representative images and quantification for Tip5 staining in Ctrl and 30 minutes and 2 hrs post compression show lack of enrichment of Tip5 in the nucleoli under compression and global significant decrease at 2 hrs. (Scale bar=10μm, N=3 independent experiments, at least n>41 per condition per experiment, with at least total of 135 nucleoli per condition ***p<0.001 in Mann-Whitney test)*

*G) Representative images and quantification for Suv4-h20 staining in Ctrl and 30 minutes and 2 hrs post compression show lack of enrichment nucleolar enrichment at 30 minutes and significant enrichment at 2 hours post compression. (Scale bar=10μm, N=3 independent experiments, at least n>48 per condition per experiment, with at least total 178 nucleoli per condition; ****p<0.0001 in Mann-Whitney test)*

### Figure S5

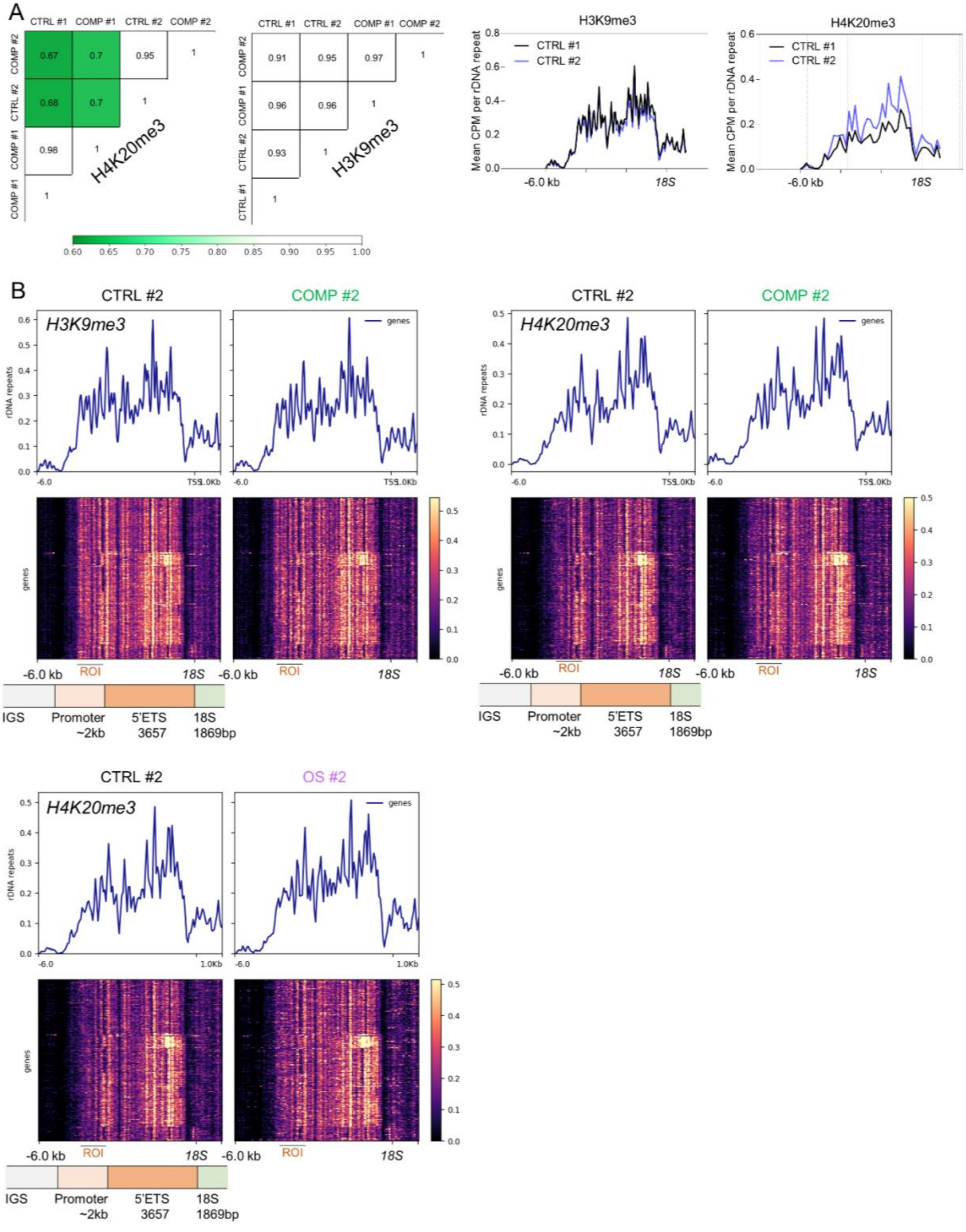
A
CTRL #1
COMP #1
CTRL #2
COMP #2
H4K20me3
H3K9me3
H3K9me3
H4K20me3
Mean CPM per rDNA repeat
CTRL #1
CTRL #2
-6.0 kb
18S
B
CTRL #2
COMP #2
OS #2
genes
rDNA repeats
-6.0
TSS
1.0Kb
ROI
IGS
Promoter
~2kb
5'ETS
3657
18S
1869bp

***Figure S5 in relation to Figure 7***

*(A) Reproducibility analysis of H3K9me3 and H4K20me3 CUT&Tag datasets mapped to rDNA loci (18S, 5.8S and 28S regions). Left, Pearson correlation heatmaps showing signal concordance between biological replicates in control (CTRL) and compression (COMP) conditions for each histone mark. Right, profiles of mean CPM-normalized coverage across rDNA regions aligned relative to the 18S rRNA start site. Profiles show CPM absolute value variation between two replicates of h4k20me3 mark but high reproducibility patterns across conditions.*

*(B) Representative locus-resolved CUT&Tag signal and repeat-wide heatmaps for H3K9me3 and H4K20me3 across individual rDNA repeats. Top, coverage tracks showing CPM normalized signal intensity across the rDNA promoter and transcribed region (-6 kb to 1kb with 18S start as reference) in representative samples (CTRL #2, COMP #2, and OS #2 where indicated). Bottom, heatmaps displaying signal distribution across individual annotated rDNA repeats aligned from the upstream intergenic region (IGS) to the 18S rRNA gene. The region of interest (ROI) used for quantitative analyses is highlighted (-4.5kb to -3.5 kb with 18S start as reference). Annotation below indicates the relative positions of the IGS, promoter (~2 kb upstream), 5'ETS (3657 bp), and 18S (1869 bp) regions.*

#### Figure S6 Gels and Westernblot membranes

**In-plane compression**

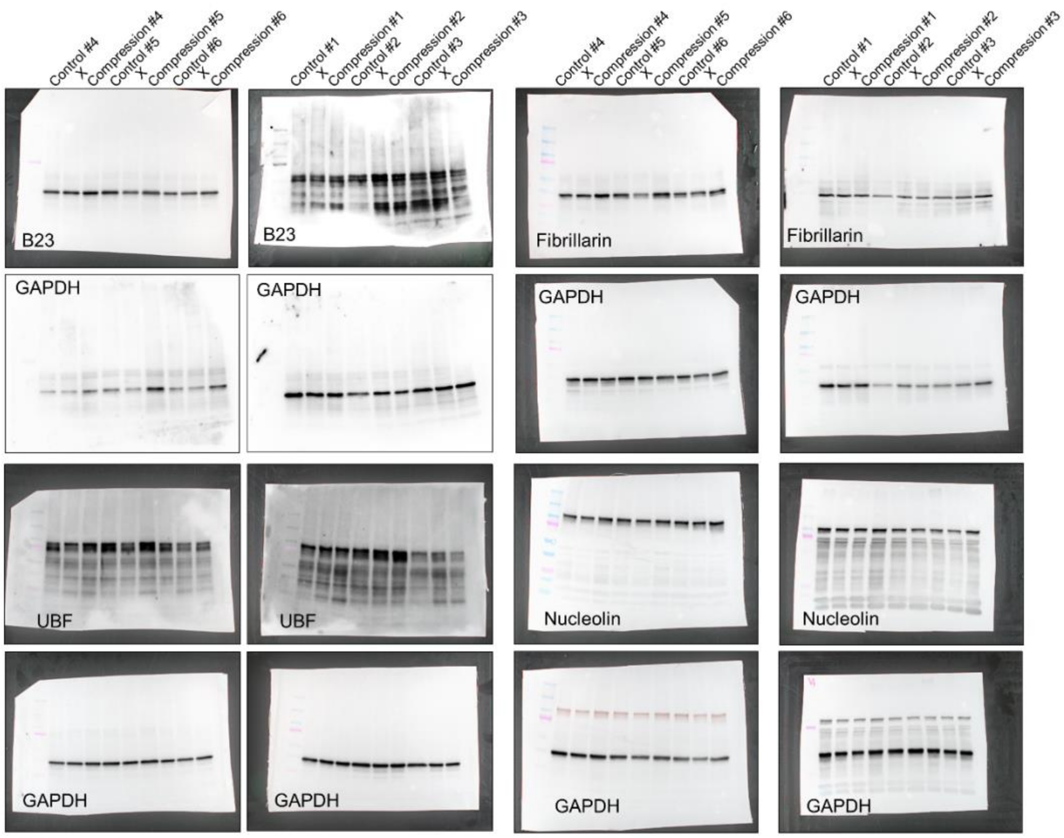


**Axial confinement**

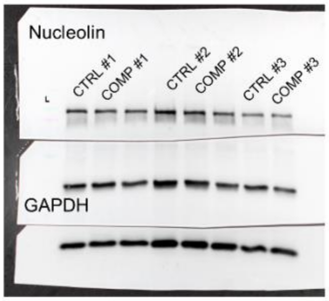

**PhosTag nucleolin**

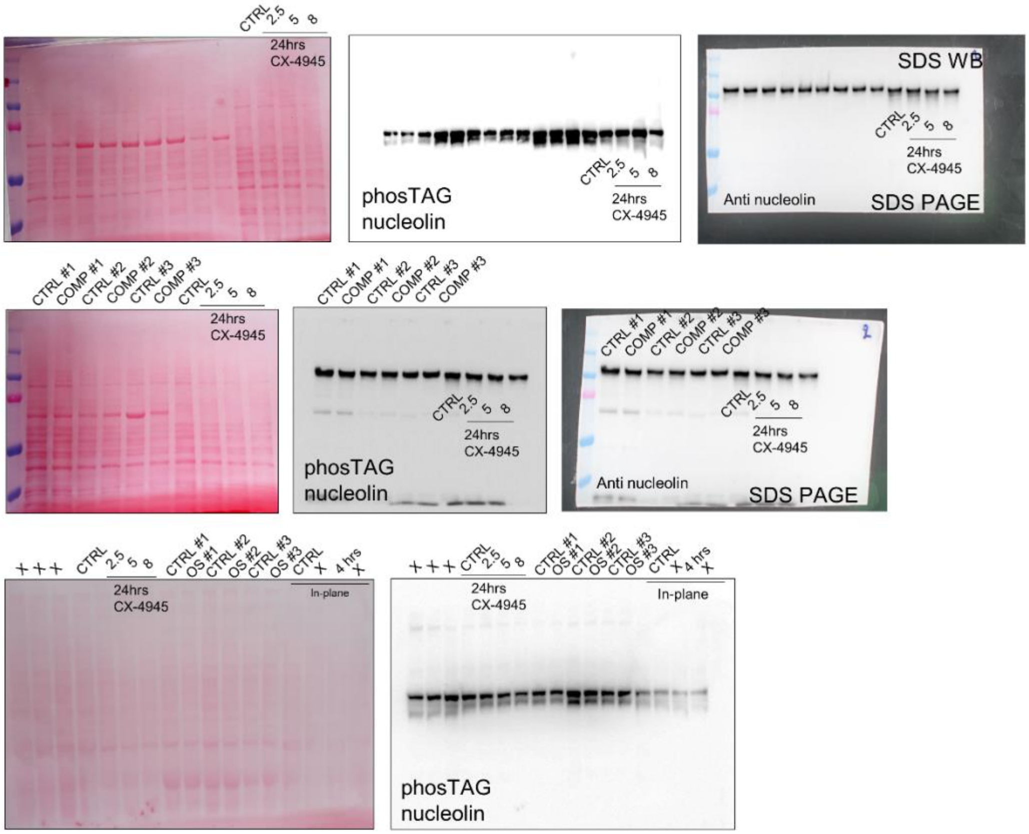


**RFP trap – Immunoprecipitation for proteomic analysis**

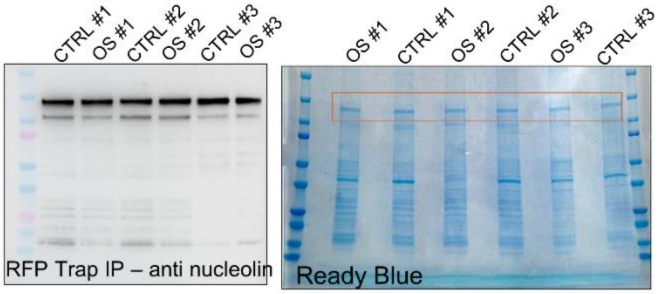


**ProQ diamond / SyproRuby**

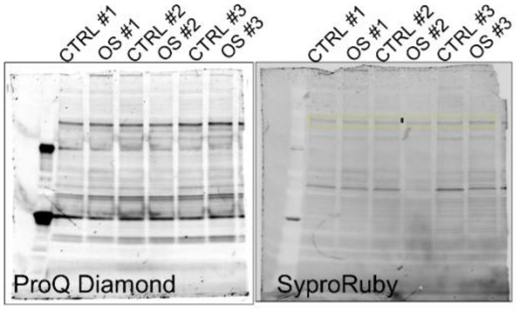
